# Native yeast kinetochore structures identify an essential inner kinetochore interaction

**DOI:** 10.64898/2026.01.30.702844

**Authors:** Mengqiu Jiang, Changkun Hu, Sabrine Hedouin, Angelica Andrade Latino, Yasuhiro Arimura, Andrew B. Stergachis, Sue Biggins

## Abstract

Kinetochores must accurately assemble on centromeres for faithful chromosome segregation. Although a conserved centromeric nucleosome is essential for kinetochore assembly, budding yeast centromeric DNA is a poor template for nucleosome formation *in vitro*, perhaps due to its intrinsic rigidity. To better understand yeast inner kinetochore assembly, we developed a one-step protocol to purify native inner kinetochore subcomplexes for structural studies. We performed cryo-electron microscopy on the purifications and generated density maps of four separate inner kinetochore complexes, two of which had not previously been visualized and may represent intermediate assembly states. We identified an Ndc10 trimerization domain that engages centromeric DNA and a pair of CBF3 complexes and is associated with substantial bending of centromeric DNA. Ndc10 trimerization is essential for kinetochore assembly and chromosome segregation. We propose that Ndc10 trimerization facilitates centromeric DNA bending to stabilize the centromeric nucleosome and inner kinetochore.

## Introduction

Faithful chromosome segregation ensures that replicated chromosomes are evenly distributed to daughter cells. Chromosome segregation is mediated by the kinetochore, a conserved protein machine that interacts with microtubules to separate chromosomes during cell division^1,2^. The kinetochore assembles on centromeres, chromosomal loci that are usually composed of thousands to millions of repetitive DNA base pairs that are not sequence-defined^3^. Centromeres are therefore epigenetically specified by specialized centromeric nucleosomes containing a histone H3 variant called CENP-A^4,5^.

Centromeric chromatin serves as a platform to assemble the inner kinetochore, which is made up of a large <u>c</u>onstitutive <u>c</u>entromere-<u>a</u>ssociated <u>n</u>etwork (CCAN)^6,7^ (Fig. 1a). The inner kinetochore serves as a bridge to the outer kinetochore that binds to microtubules and contains multiple copies of the Mis12, KNL1 and Ndc80 subcomplexes, as well as additional microtubule associated proteins such as the Ska1 complex or its budding yeast ortholog, the Dam1 complex^8–10^. Although kinetochore components are well defined, a complete structural understanding of how these proteins assemble in the native context to make functional kinetochores is lacking.

**Figure 1.**
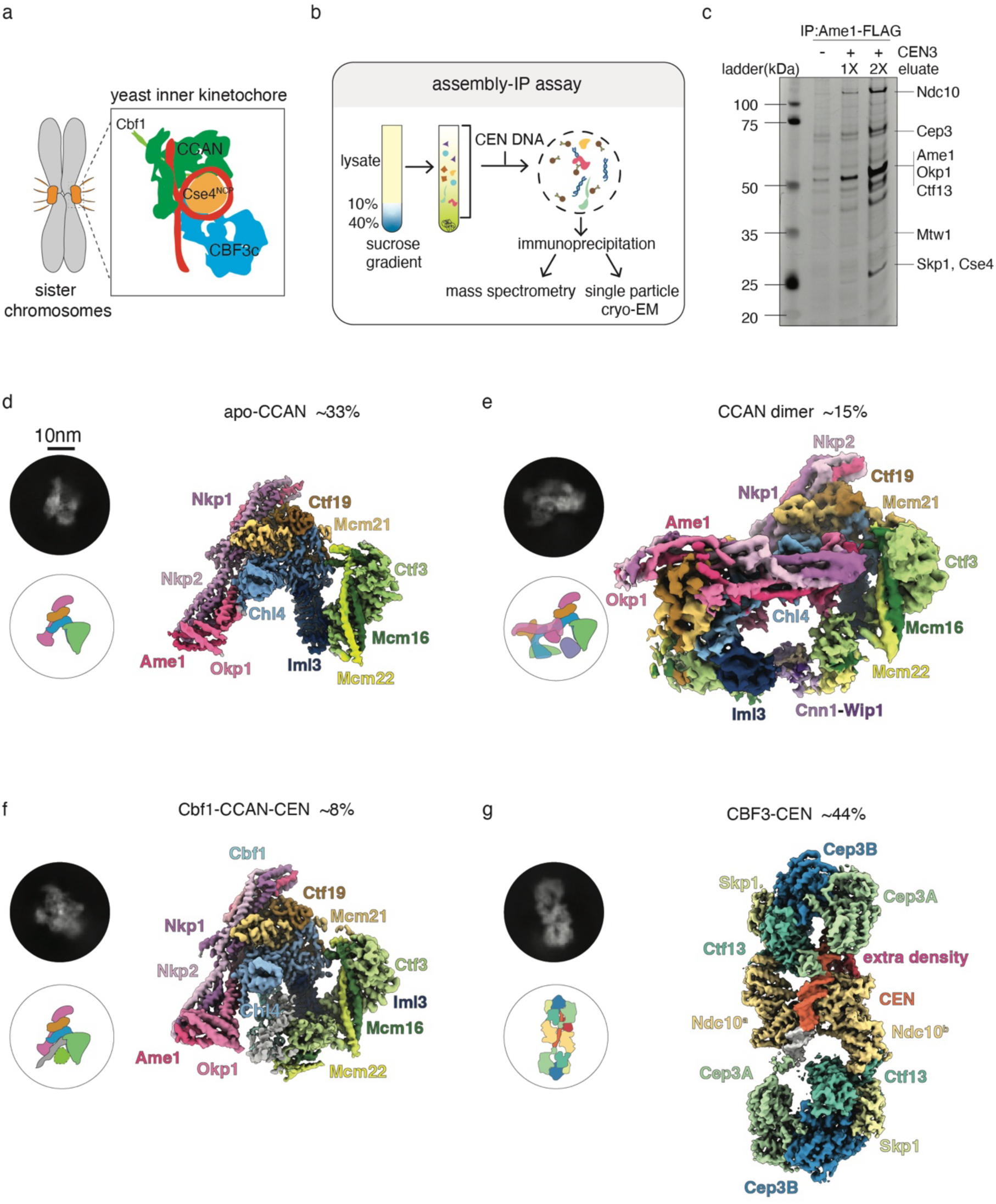
The purification of native kinetochore complexes and cryoEM overview of identified structures. a). A cartoon of a pair of budding yeast sister chromosomes with kinetochores colored in orange. The composition of the inner kinetochore complexes is illustrated in the box. CCAN is the constitutive centromere associated network, and NCP is the nucleosome core particle. b). Flowchart of assembly-IP method for native kinetochore complex purification. c). Silver-stained SDS-PAGE of Ame1-3XFLAG (SBY21782) IP or assembly-IP eluates. The presence of CEN3 DNA is indicated and the sample loading in lane 4 is twice the amount of sample in lane 3. Kinetochore proteins are labeled based on the protein band molecular weight. d-g). The four structures solved from the assembly-IP assay. Representative 2D classifications and the density maps are shown for each structure. The name of the proteins are labeled in the corresponding color and the relative percentage of each complex detected is reported.

In *Saccharomyces cerevisiae*, the centromere is unique and sequence-specified by a ∼125 bp sequence that assembles a single essential centromeric nucleosome required for kinetochore assembly^11,12^. The yeast centromere is divided into three <u>c</u>entromere <u>d</u>etermining <u>e</u>lements: CDEI, CDEII and CDEIII^3,13^. The CDEI and CDEIII elements are sequence defined^14^: CDEI is a palindromic, conserved DNA sequence that interacts specifically with a dimer of the transcription factor Cbf1^15,16^, while CDEIII is a 25 bp element containing several key nucleotides that are responsible for the recruitment of the CBF3 complex^17–19^. In contrast, CDEII is not sequence defined, but it is extremely AT-rich, with many homopolymer runs of A or T repeats spanning ∼80 bp that are required for accurate centromere function^20^. However, A-T homopolymers are relatively stiff and are predicted to be poor templates for nucleosome formation^20^. Consistent with this, reconstitution experiments in which Cse4, the yeast CENP-A histone variant, were added to native CEN DNA result in limited and unstable nucleosome formation^21,22^. Thus, although the yeast centromere must form an essential centromeric nucleosome, it is paradoxically a very poor template for nucleosome formation. To address this problem, most inner kinetochore reconstitutions have replaced CDEII with a strong nucleosome positioning sequence or used a stabilizing antibody that blocks binding of the essential CCAN component Mif2 to assemble stable centromeric nucleosomes^21–24^. While chaperone-mediated loading at centromeres may create a high local concentration of Cse4 that could overcome the intrinsic stiffness of CDEII in cells, additional mechanisms are required to stabilize the centromeric nucleosome. For example, we recently found that the CCAN subcomplex OA (Okp1-Ame1) ensures centromeric nucleosome stability^25^, and it is likely there are additional unidentified mechanisms that stabilize the centromeric nucleosome.

Yeast kinetochore assembly is initiated by the essential CBF3 complex (CBF3c) that enables Cse4 deposition^2^. The CBF3core contains a Cep3 dimer, Ctf13, and Skp1. The Ndc10 protein loosely binds to the CBF3core to form the full CBF3c^26^. CBF3c interacts with the CDEIII element via a CCG motif that binds to the GAL4-like domain of Cep3^27–30^. Although it was proposed that CBF3c helps bend DNA to contribute to nucleosome formation^26,28,29^, it is not clear whether nor how CBF3c bends DNA. The behavior of Ndc10 at the centromere is also unclear. Ndc10 is localized to kinetochores throughout the cell cycle *in vivo*^31^, but structural studies have suggested that Ndc10 must dissociate from CBF3c after centromeric nucleosome formation^21,32^.

High-resolution structures of recombinant budding yeast inner kinetochore complexes, including CBF3–CEN assemblies^32,33^, apo-CCAN^34^, and CCAN bound to a Cse4 nucleosome^21,23^, have provided a detailed framework for how core components can engage centromeric DNA and one another. To achieve stability and homogeneity, these studies typically used engineered constructs, such as modified CDEII sequences or stabilizing antibodies^21,22^, and in some cases truncated or modified proteins^32^, which may limit information about native DNA bending, post-translational modifications, and transient assembly states. We build on this work by analyzing inner kinetochore complexes assembled from native budding yeast components on CEN DNA, with the goal of identifying conformations and intermediate states that are challenging to access in fully reconstituted systems.

We developed a method to assemble and efficiently enrich native budding yeast kinetochore complexes for cryoEM studies and identified native structures that largely agree with previous work but contain previously unknown features. We observed significant DNA bending in the Cbf1-CCAN-CEN and CBF3-CEN complexes at the CDEI–CDEII and CDEII–CDEIII junctions. Specifically, the bend at the CDEII–CDEIII junction correlates with an additional structural component, not observed in earlier reconstitutions, that we identify as an Ndc10 trimer. Consistent with an important role for Ndc10 trimerization in inner kinetochore assembly, mutants that abolish the trimer are inviable and exhibit severe defects in kinetochore formation. Together, our work uncovers structural details that contribute to understanding how the yeast inner kinetochore assembles and lay the foundation for future studies to elucidate kinetochore function.

## Results

### Purified *de novo* assembled-IP kinetochores consist of many states

Kinetochore proteins are expressed at low levels and a single centromeric nucleosome assembles the entire yeast kinetochore, so native material for structural analyses is limited^9^. We previously isolated native kinetochore material from *S. cerevisiae* using a 3xFLAG epitope tag on the Dsn1 kinetochore protein^35^. While this method is useful for biochemical and biophysical assays^35–37^, cryoEM is not possible due to contaminating proteins and aggregates and the prep cannot be purified through gel filtration steps due to its large size and small quantity. In addition, the Dsn1 purification is limiting for inner kinetochore proteins^35^. To address these issues, we developed a protocol to obtain native kinetochores that have fewer non-specific co-purifying proteins and a higher yield of the inner kinetochore. We arrested budding yeast cells in mitosis with the microtubule destabilizing drug benomyl and generated cell lysates. We then pre-cleared the lysates via a sucrose gradient to decrease contaminating proteins (Supplementary Fig. 1a). We next immunoprecipitated (IP) three different inner kinetochore proteins: Ame1, Chl4 and Mif2, and found that an Ame1 IP best enriches for inner kinetochore proteins (Supplementary Fig. 1b).

To further recruit inner kinetochore complexes, we combined a well-established kinetochore assembly assay^38^ with the Ame1-3xFLAG immunoprecipitation. We added 500 bp centromeric DNA fragments containing 125 bp of CEN3 and additional flanking pericentromeric DNA on both ends into the clarified lysate and let kinetochores assemble for 90 minutes. We then immunoprecipitated Ame1-3xFLAG and eluted the material from beads using FLAG peptide. We call this purification method an “assembly-IP” (Fig.1b). When the assembly-IP was assayed by silver-stain analysis of SDS-PAGE and immunoblotting against kinetochore proteins (Fig. 1c and Supplementary Fig.1c), we observed enrichment of inner kinetochore proteins such as the CBF3 and Ame1-Okp1 complexes. To fully analyze the composition of the final assembly-IP product, we performed mass spectrometry. We found that most kinetochore proteins were present and the inner kinetochore proteins were detected with relatively high peptide-spectrum matches (PSMs)(Table S1). Taken together, these data showed that the assembly-IP method enriches for native kinetochore material containing the inner kinetochore.

We next analyzed both the Ame1-3xFLAG IP and the assembly-IP eluates by cryoEM (Supplementary Fig. 1d-g, Supplementary Fig. 2) and compared the corresponding structures. When centromeric DNA was added prior to the IP, we identified four kinetochore complexes at reasonable resolution (3.3 Å to 6.5 Å). We built models for all four complexes by using either previously solved related kinetochore subcomplex structures as the initial models^21,33,34^ or AlphaFold predictions^39^. In summary, we identified a single copy apo-CCAN (33% of the particles at 3.3 Å resolution, Fig. 1d), a CCAN dimer structure in a different conformation compared to a prior reconstituted version (15% at 6.5 Å resolution, Fig. 1e), a Cbf1-CCAN-CEN complex (8% at 3.9 Å resolution, Fig. 1f), and a CBF3-CEN complex with a significant extra structural density that was not visualized in a previous study (44% at 3.5 Å resolution, Fig. 1g and Supplementary Fig. 2). It is noteworthy that even though we observed a 2D classification corresponding to a nucleosome complex (Supplementary Fig. 1e, pink box), we could not generate a density map for that structure. This observation suggests that the nucleosome is not stable upon cryoEM sample preparation, consistent with previous studies^21,22^.

### *De novo* assembled structures adopt unique conformations

Although the *de novo* assembled structures we identified are overall similar to previously reconstituted complexes, some reveal features that were not apparent in earlier work. The apo-CCAN complex we obtained is highly similar to previously solved recombinant CCAN structures^34^, so we focused our analysis on three additional complexes we observed when CEN DNA was added prior to the purification. One corresponds to a Cbf1–CCAN–CEN assembly in which we positioned the Cbf1 dimer on CDEI according to prior crosslinking and reconstituted structures. While the Cbf1-CCAN interaction is the same as previous reconstituted structures^16,21^, the CEN DNA bends roughly 15° more toward Cnn1–Wip1 than the reconstituted complex containing a modified CDEII sequence^21^ (Supplementary Fig. 3a-b). This increased curvature is compatible with the intrinsic bending reported for the native, AT-rich CDEII in biochemical assays^40^.

Another structure we detected corresponds to a CCAN dimer whose abundance increased with added CEN DNA, indicating a DNA-dependent CCAN–CCAN association (Fig. 2a). The dimer adopts a more open conformation than previously described recombinant CCAN dimers^34^ (Fig. 2b), with extra density connecting the two complexes, but the local resolution is insufficient for unambiguous assignment. We therefore consider this CCAN dimer a candidate DNA-dependent inner kinetochore state, although it is difficult to determine whether it represents a physiological assembly intermediate. An AlphaFold 3 model suggests that the flexible density could arise from a Mif2 segment contacting Okp1^39^ (Supplementary Fig. 3e-f). We also predicted that Okp1 from CCAN interacts with Chl4 from CCAN^2^ through an electrostatic interaction (Supplementary Fig. 3g). Disrupting this interaction resulted in synthetic lethality in combination with the *mif2-3* mutant at a semi-permissive temperature (Supplementary Fig. 3h-i), consistent with the CCAN dimer facilitating inner kinetochore function *in vivo*.

**Figure 2.**
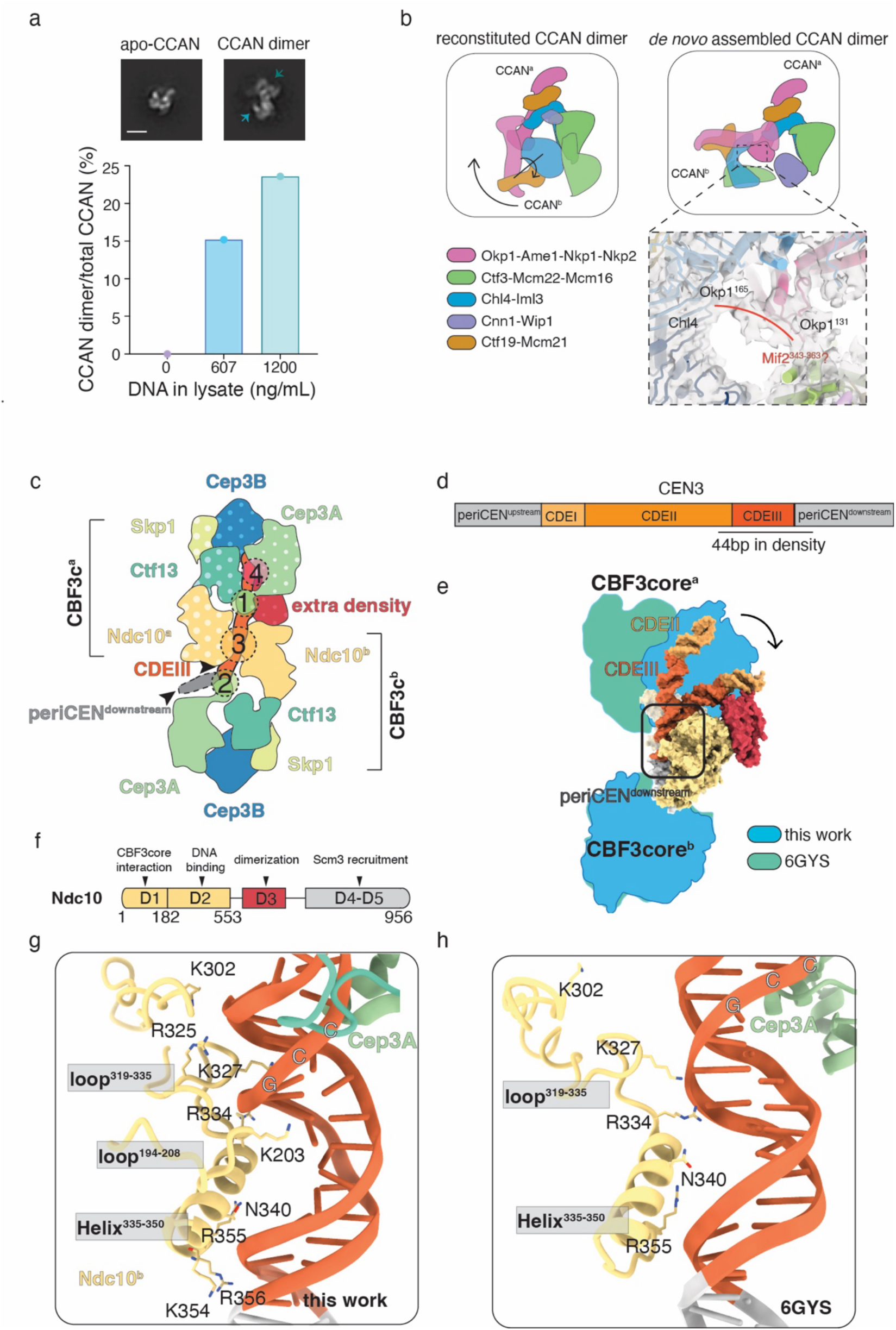
Comparison of native and recombinant reconstituted CBF3-CEN structures. a). Representative 2D averages of the apo-CCAN and CCAN dimer from negative stain EM (scale bar=10 nm). Graph depicts the ratio of CCAN dimer to total CCAN detected when 0, 607 or 1200 ng CEN DNA was added per ml of precleared lysate. b). Comparison between a recombinant reconstituted CCAN dimer^23,34^ (left upper panel) and the native CCAN dimer (left bottom panel). CCAN^a^ is shown in solid colors and CCAN^b^ is shown in transparent colors. Different colors represent different protein subcomplexes as indicated with color blocks. The symmetry axis of the reconstituted CCAN dimer is noted. In the native CCAN dimer, the CCAN^b^ will move toward CCAN^a^ and turn. Zoomed-in density is shown on the right panel to show the extra density that connects Okp1 from CCAN^a^ to Chl4 from CCAN^b^. Red line traces the putative Mif2^343–363^ density. c).Cartoon of the organization of the CBF3-CEN complex. Each CBF3c protein is colored and labeled. CBF3c^a^ are colored in polka dot and CBF3c^b^ is colored in solid colors. The arrowheads point to CDEIII that tightly interacts with CBF3c and periCEN^downstream^. The four protein-DNA interaction surfaces are outlined by dashed circles and labeled with numbers. d). Color scheme of CEN3 used in assembly assay. Based on our structure, 44 bp DNA protected by CBF3 complexes are underlined. The periCENs are colored in grey. CDEI is colored in light orange. CDEII is colored in orange. CDEIII is colored in dark orange. e). Illustration of CEN DNA bending by superimposing the recombinant reconstituted CBF3-CEN^33^ (PDB: 6GYS) and native CBF3-CEN structures on CBF3core^b^. The movement of DNA is indicated by the arrow. The extra density is colored in bright red. The interaction surface between Ndc10^b^ and CEN DNA is boxed with a black rectangle, and the details are highlighted in (g.) and (h.). To illustrate that CEN DNA bending corresponds to the angle in a Cse4 nucleosome, a previously solved Cse4 nucleosome (PDB:7k78) structure^22^ is shown. The CEN DNA is colored in the same scheme as on the left. Cse4 is colored in cyan. PeriCEN, CDEIII and the small part of CDEII are in transparent shade to compare the bending angle with the CEN DNA on the left. f). Cartoon of Ndc10 domain organization. D1-D2 are labeled and colored in yellow, which is well-structured and resolved in our structure. D3 is colored in bright red, and its structure is solved in this work. D4-D5 are not resolvable and colored in grey. The proposed function of each domain is labeled. g-h).Interaction between Ndc10^b^ and CEN DNA in the native CBF3-CEN structure (g) and the recombinant reconstituted structure ((h),PDB: 6GYS) shows that there are more Ndc10^b^ residues in the native structure that contact the CEN DNA. Ndc10^b^ is colored in yellow, the CDEIII DNA is colored in orange and GAL4 from Cep3A is colored in light green. The CCG motif is highlighted.

### The native CBF3 complexes interact with CEN DNA tightly and induce a sharp bend

The final structure we identified is a CBF3c dimer containing a previously undetected extra density that interacts with CEN DNA. Using AlphaFold 3 prediction and referring to a previously published reconstituted CBF3-CEN structure^33^, we were able to generate a native CBF3-CEN complex model. The CEN DNA was threaded between the two CBF3 complexes and contacted them in four places (Fig. 2c-d). Two of the interactions with CEN DNA were previously reported^29,32,33,41^ and represent Cep3A from CBF3c^a^ binding to the conserved CCG motif in CDEIII and Cep3A from CBF3c^b^ binding to periCEN DNA through a non-sequence specific interaction (Fig. 2c, interactions 1 and 2 and Supplementary Fig. 4a). The third CEN DNA interaction is the Ndc10 proteins from each CBF3c clamping onto CDEIII and the periCEN DNA through non-specific DNA interactions^29,33^ (Fig. 2c, interaction 3). Finally, we observed an interaction between the extra density with the CEN CDEII DNA that is adjacent to the CDEIII element (Fig. 2c, interaction 4).

To compare the native and reconstituted structures, we superimposed the two CBF3core^b^ complexes from each structure. When they are aligned, we found that the CBF3c^a^ complexes bind to CEN DNA differently, which results in a large bend and kink in the CDEIII element close to the DNA downstream of the centromere (periCEN^downstream^) in the structure we identified (Fig. 2d-e, and movie 1). The bend we identified matches the angle of the DNA in a reconstitution of the centromeric nucleosome (Fig. 2e), suggesting that CBF3 binding can stabilize a DNA trajectory that is geometrically compatible with formation of the centromeric nucleosome^32^.

We also detected more Ndc10 residues interacting with CEN DNA in the native structure (Fig. 2g) compared to the reconstituted one (Fig. 2h). The Ndc10 polypeptide sequence is roughly divided into five domains with D2 mediating DNA binding^42^ (Fig. 2f). Consistent with prior work^33^, D2 domain residues in Ndc10^b^ insert into the major groove immediately downstream of the CCG motif that interacts with Cep3A. This reinforces the Cep3A interaction with the CEN DNA and positions Ndc10 deeper in the major groove (Fig. 2g). We also detected additional CEN DNA contact points through two lysines from Ndc10^b^ loop^319–335^ and another Ndc10^b^ loop^194–208^ that interacts with the DNA sugar backbone (Fig. 2g-h). Ndc10^a^, the other copy of Ndc10, interacts with the 3’ end of CDEIII and some periCEN^downstream^ through the same residues. Together with the strong extra density that contacts the CDEII–CDEIII junction, these features indicate that Ndc10 can stabilize a sharply bent DNA conformation at CDEIII that is compatible with our overall CBF3–CEN architecture.

### Identification of an Ndc10 trimer domain

The extra structural density that closely interacts with Ndc10^b^ and several base pairs at the end of CDEII is very well-resolved and present in all of our CBF3-CEN 3D classifications and even in the structure of the monomeric CBF3 core and CEN DNA, indicating a strong interaction between the extra density and Ndc10, the core CBF3 complex and the centromeric DNA (Supplementary Fig. 4b, class 6). In addition, even though the CBF3-CEN structure is symmetrical in the native structure, only CBF3^a^ interacts with the extra density and the other side of the CBF3 complex lacks a comparable feature. Taken together, these data suggest that the extra density represents an early step in inner kinetochore assembly that may have an important functional role, such as stabilizing the centromeric nucleosome.

To identify the extra density, we performed local 3D classification by masking particles with the strongest extra density in cryoSPARC^43^ and then performing local refinement using the extra density center as the fulcrum. This improved the local resolution and density connectivity, allowing the side chains to become clear (Supplementary Fig. 4c-d). We then predicted its sequence through model-angelo^44^ and ran it through a blast prediction program. The extra density was predicted to correspond to three chains of helix-loop-helix (HLH) hairpin structures that all have about 30% sequence identity with Ndc10 residues 579-639 (Supplementary Fig. 4e). When we modeled the extra density with the Ndc10^579–639^ sequence, the overall fit was good, particularly for some bulky side chains of phenylalanine and tryptophan (Supplementary Fig. 4d). In this model, two pairs of Lys610 and Lys611 from two Ndc10^579–639^ segments contact the DNA, consistent with the strong density at the CDEII–CDEIII junction (Fig. 3a).

**Figure 3.**
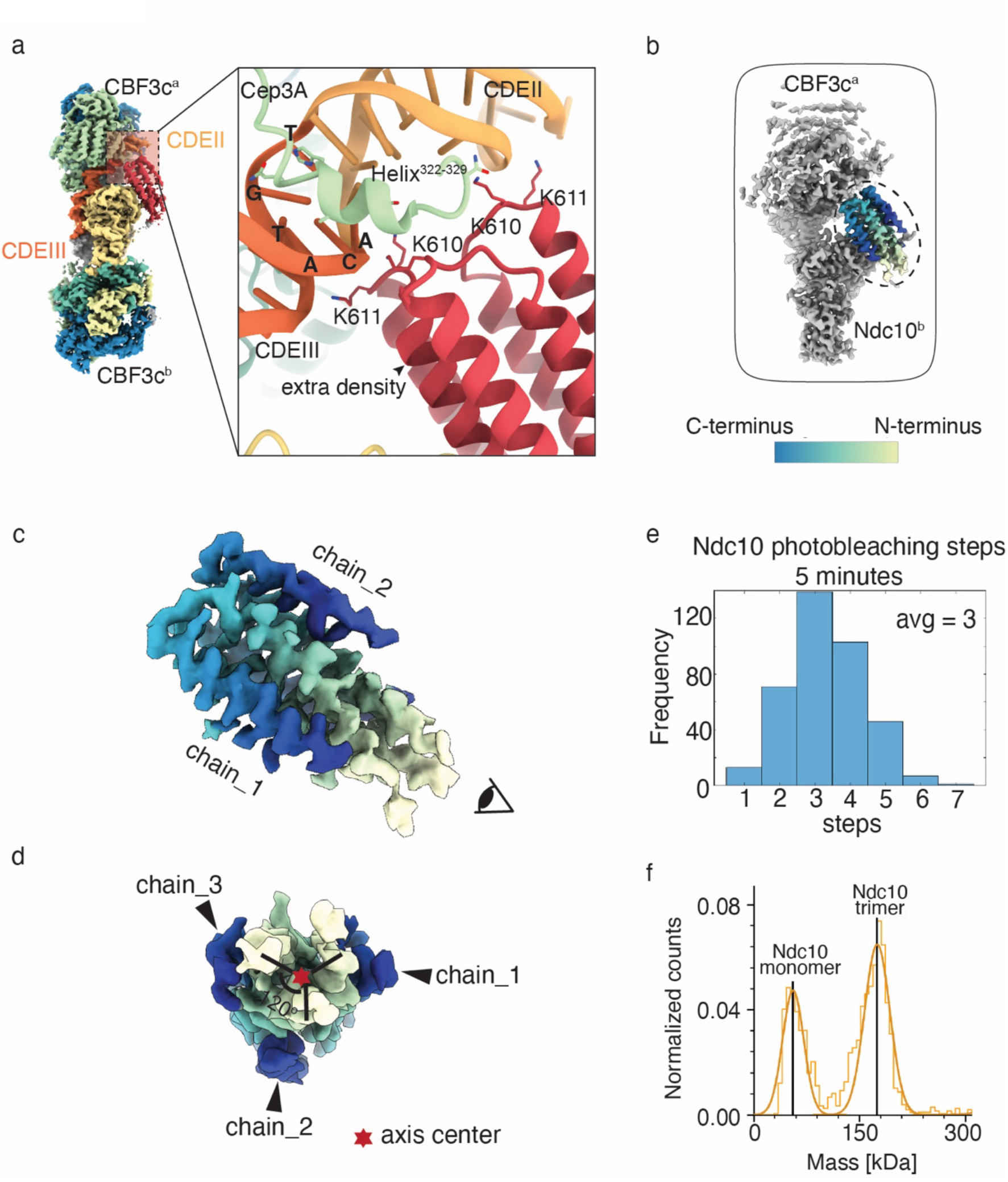
The extra structural density in the native CBF3-CEN complex is a trimer of Ndc10^579–639^. a). Detailed interaction between the extra density and the CEN DNA. The model of the extra density is colored in bright red. CDEIII is in dark orange and CDEII is in orange. b). Focused refinement of the density map shows well-resolved extra density in the box. The extra density is colored and circled with a dashed line. The C-terminus of the helix-loop-helix is colored in blue, and the N-terminus of the helix-loop-helix is colored in pale yellow. c). Zoom-in view of the extra density adopting the same view from b. d). Zoom-in view of the extra density observed from the cartoon eye angle in (c). Each of the chains is labeled and the symmetry axis is marked by a red star. The C-terminus and N-terminus are indicated by blue and pale yellow, respectively. e). The frequency of photobleaching steps on kinetochores assembled from extracts from Ndc10-GFP cells (SBY22191) via a single molecule TIRF assay after *de novo* assembly for 5 minutes. f). Mass photometry of Ndc10^D3-D5^ purified from *E.coli*. Two major peaks were observed and correspond to an Ndc10^D3-D5^ monomer and trimer, respectively.

It was previously proposed that the *K. lactis* Ndc10 forms dimers through D3^42^. In our structure, residues 579–634 within the D3 region mediate a threefold symmetric oligomer, so we refer to this module as the Ndc10 D3 trimer (Fig. 3b–d). To validate that Ndc10 forms a trimer, we used a TIRF microscopy assay to analyze the Ndc10 copy number on assembled kinetochores via photobleaching^45,46^. In this assay, fluorescently labeled CEN DNA molecules were tethered to a coverslip and incubated with lysate made from cells expressing Ndc10-GFP. Within 5 minutes of assembly, there were approximately 3 steps of Ndc10-GFP photobleaching for 40% of the CEN DNA molecules (Fig. 3e), consistent with an Ndc10 trimer. Our prior work indicated that the inner kinetochore takes more than 30 minutes to assemble in this assay^45^, suggesting that the Ndc10 trimer forms in the absence of the rest of the inner kinetochore. The number of Ndc10 proteins stayed relatively similar after 30 minutes of assembly (Supplementary Fig. 4f), indicating that three Ndc10 molecules are maintained as the inner kinetochore is assembled.

To further validate Ndc10 trimerization, we purified a recombinant Ndc10 fragment spanning residues 541 to 956 (D3-D5) from bacteria (∼52kDa). Mass photometry of this fragment revealed two major species corresponding to a monomer (∼53kD) and a trimer (∼150kD)(Fig. 3f), consistent with D3-mediated trimer formation in solution. An AlphaFold 3^39^ model of Ndc10^D3-D5^ (Supplementary Fig. 4g, upper panel) predicts a trimeric arrangement that closely matches the cryoEM density, including the relative orientation of helices and positions of bulky side chains (Supplementary Fig. 4g, lower panel).

### Docking of the Ndc10 trimer on CBF3 is important for kinetochore assembly

The Ndc10 trimer docks onto CBF3 through an interaction with the D1-D2 domains of the Ndc10^b^ protein. From our CBF3-CEN model, we identified six residues within the D1-D2 domain of Ndc10 (S493, D494, S497, D501, H505 and K507) that contact the D3 trimer density (Fig. 4a). To determine whether this interface contributes to kinetochore function, we mutated these residues to alanine (*ndc10^6A^*). The *ndc10^6A^* mutant showed no obvious growth defect on its own but was lethal with the *mif2-3* inner kinetochore mutant at a semi-permissive temperature, indicating that this surface becomes critical when inner kinetochore function is compromised (Fig. 4b). To further explore the effects of the *ndc10^6A^* mutant, we performed a *de novo* kinetochore assembly assay^38^. Ndc10^WT^ and Ndc10^6A^ assembled onto CEN DNA normally (Supplementary Fig. 4h). In contrast, assembly of Mif2 and Okp1 was strongly reduced and Cse4 recruitment was modestly impaired in the *ndc10^6A^*lysate, consistent with a defect in inner kinetochore assembly rather than an initial CBF3–DNA binding (Supplementary Fig. 4h). At the single-molecule level, TIRF-based assembly assays with Mif2-GFP similarly showed reduced Mif2 co-localization with CEN3 DNA in *ndc10^6A^* extracts (Fig. 4c). Taken together, these data support a model in which the interaction surface between the Ndc10 D3 trimer and the D1/D2 region of the Ndc10^b^ protein contributes to efficient inner kinetochore assembly.

**Figure 4.**
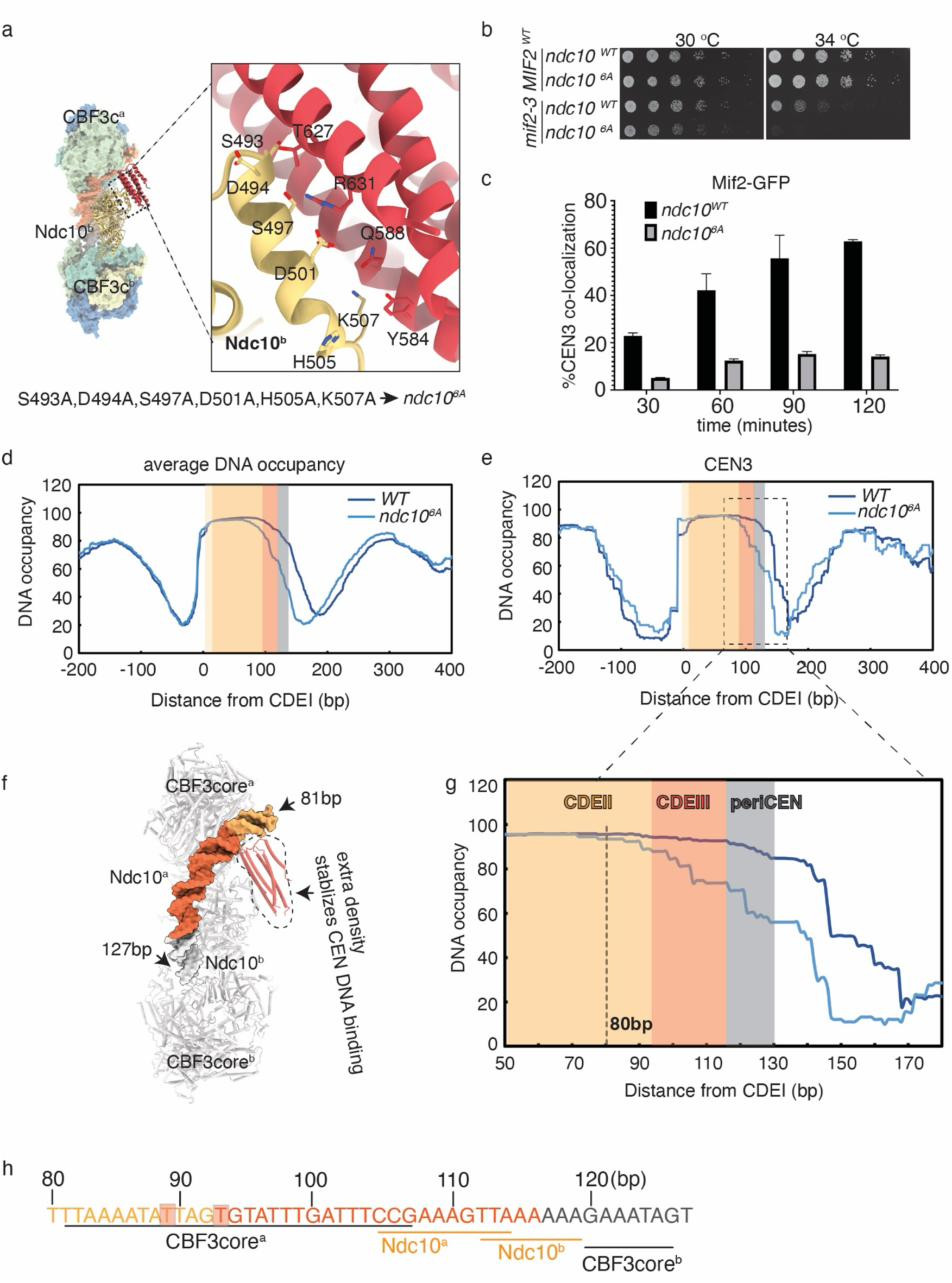
The interaction between Ndc10^D3_trimer^ and Ndc10^D1−D2^ from Ndc10^b^ stabilizes the association of the kinetochore and centromere. a). Residues involved in the interactions between Ndc10^D3_trimer^ and Ndc10^b^. Ndc10^b^ is yellow and the Ndc10^D3_trimer^ is bright red. The residues that are mutated in *ndc10^6A^* are labeled. b). Five-fold serial dilutions of WT (SBY3), *ndc10^6A^* (SBY23670), *mif2-3* (SBY21479) and *ndc10^6A^ mif2-3* (SBY23913) cells were plated on 2% glucose at 30 °C and 34 °C. c). Co-localization of Mif2-GFP and CEN3 DNA measured by a single molecule TIRF assay using *MIF2-GFP* (SBY22094) and *MIF2-GFP ndc10^6A^* (SBY24238) strains. d). Average DNA occupancy of all centromeres in WT (SBY3) or *ndc10^6A^* (SBY23670) yeast strains. WT or *ndc10^6A^* strains were traced with dark blue or light blue lines, respectively. The beginning of CDEI is assigned as 0 bp. CDEI is pale yellow, CDEII is orange, CDEIII is dark orange and the periCEN is grey. e). DNA occupancy of CEN3 and its neighboring DNA in WT or *ndc10^6A^* strains were traced with dark blue or light blue lines, respectively. The beginning of CDEI is assigned as 0 bp. CDEI is pale orange, CDEII is orange, CDEIII is dark orange and the periCEN is grey. f). Surface showing relative direction of CEN DNA bound to CBF3. g). Zoom-in view from f. The significant drop of DNA occupancy starting in CDEII 80 bp from CDEIII is highlighted. h). The sequence of CEN DNA that is protected by the CBF3-CEN complex is illustrated with CDEII in light orange, CDEIII in dark orange and the periCEN in grey. The regions of CEN DNA that interact with components of the CBF3 complex are labeled. The two bases that are boxed in a pink shade interact with extra density.

The presence of the Ndc10 D3 trimer correlates with stronger and more continuous DNA density at the CDEII–CDEIII junction than in a reconstituted CBF3core-Ndc10(D1–D2) structure, suggesting that inclusion of the D3 trimer enhances the stability or occupancy of CBF3–DNA contacts. To test this *in vivo*, we analyzed DNA occupancy at endogenous centromeres in WT and *ndc10^6A^* cells using single molecule chromatin fiber sequencing (Fiber-seq)^47^. Spheroplasted cells were exposed to an N6-adenine DNA methyltransferase (m6AMTase) that nonspecifically methylates accessible regions of DNA. The resulting m6A-labeled chromatin fibers were then analyzed using long-read single-molecule sequencing at greater than 100× genomic coverage, enabling detection of individual protein-binding events with nearly single-nucleotide resolution at all 16 centromeres.

Using computational tools, we calculated the percentage of DNA occupancy at each bp surrounding each centromere. Roughly 160 bp DNA is protected at centromeres in wild-type cells, encompassing ∼10 bp upstream of the CDEI element, the entire ∼120 bp of CEN DNA, and ∼30 bp downstream of the CDEIII element, corresponding to the binding of Cbf1, the Cse4 nucleosome and the CBF3 complex as previously reported^25^ (Fig. 4d and 4e, dark blue line). Strikingly, the nucleosome footprint at centromeres was shorter in the *ndc10^6A^* cells, occupying less DNA predominantly on the CDEIII and CDEIII-adjacent sides (Fig. 4d and 4e, light blue line). At individual centromeres such as CEN3, the reduction in occupancy begins ∼80 bp into CDEII and extends through the CDEII–CDEIII junction, overlapping the region that engages CBF3 and Ndc10 D3 in our native structure (Fig. 4f–h). Together, these observations indicate that the *ndc10^6A^* mutation decreases DNA protection over the CDEII–CDEIII region *in vivo*, consistent with a role for the interface between the Ndc10 D3 trimer and the D1/D2 domain in stabilizing CBF3–DNA complexes at centromeres.

### Mutating the Ndc10^D3_trimer^ self-interaction surface abolished Ndc10 function *in vivo*

The trimerization of Ndc10 depends on the hydrophobic interface formed by the three chains of Ndc10 D3. To test the effects of disrupting the trimer, we mutated residues F593, F596, F600, and F604, which pack the three α-helices tightly together (Fig. 5a). We mutated the four residues to alanine or aspartic acid (*ndc10 ^trimer_A^* or *ndc10 ^trimer_E^*). Mass photometry of recombinant Ndc10 D3-D5 fragments carrying either mutation showed loss of the ∼150 kDa trimer peak and the presence of a single major species at ∼55 kDa, indicating that both mutants disrupt D3-mediated trimerization *in vitro* (Supplementary Fig. 5a,b). We next expressed the mutants in a strain where the endogenous Ndc10 protein was fused to an auxin-inducible tag to control its degradation (*ndc10-AID*)^48^. As expected, the *ndc10-AID* strain dies on auxin and is rescued by a wild-type copy of *NDC10* (Fig. 5b). Strikingly, the two *ndc10* mutants were inviable on auxin, indicating that the hydrophobic interactions that stabilize the D3 trimer are essential for Ndc10 function *in vivo* (Fig. 5b).

**Figure 5.**
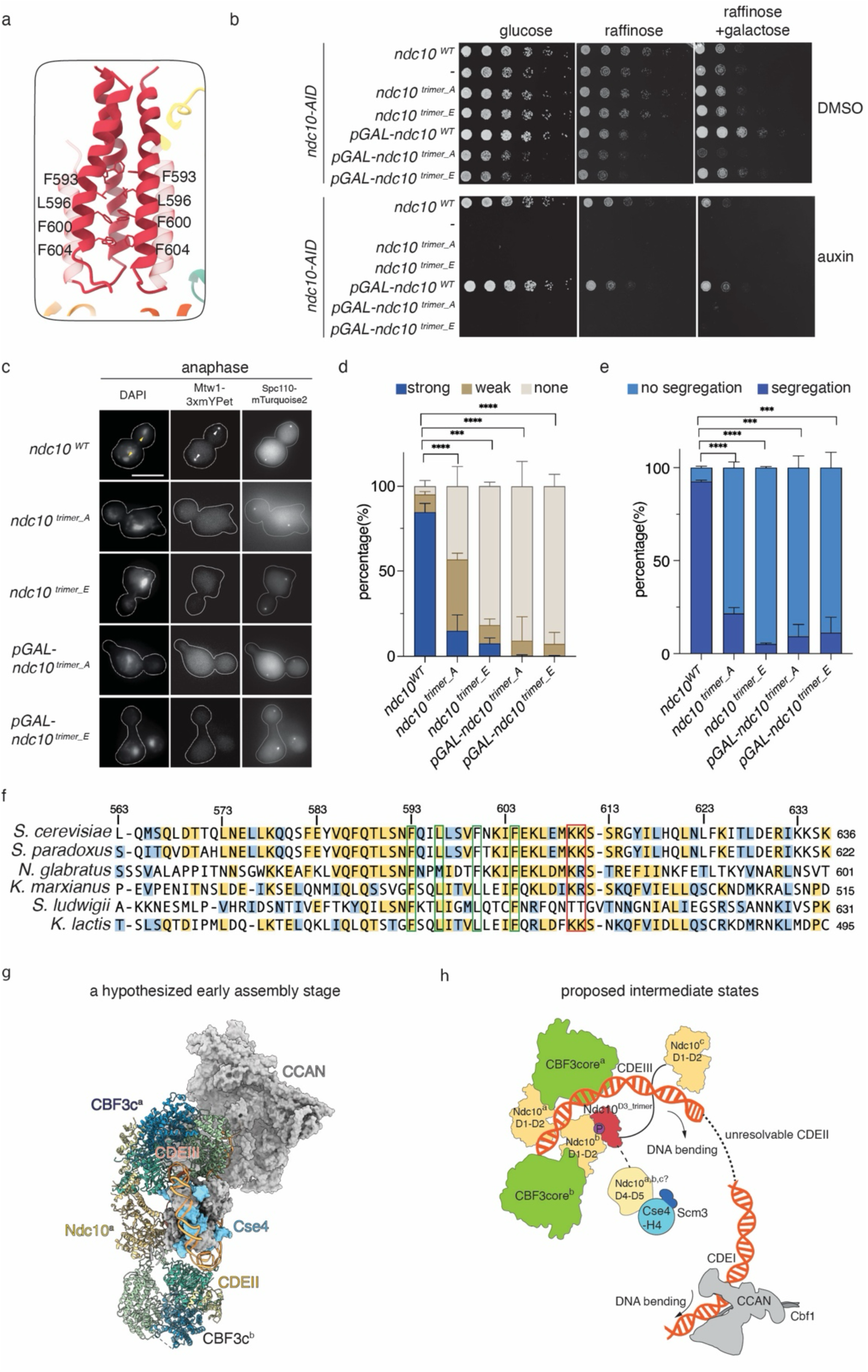
Ndc10 trimerization is essential for kinetochore assembly and cell viability. a). Zoom-in model shows residues involved in the Ndc10 trimerization. b). Five-fold serial dilutions of WT (SBY12796), *ndc10-AID* (SBY21327), *ndc10-AID ndc10^trimer_A^* (SBY24533), *ndc10-AID ndc10 ^trimer_E^*(SBY24478), *ndc10-AID pGAL-ndc10^WT^* (SBY24690), *ndc10-AID pGAL-ndc10^trimer_A^* (SBY24689), *ndc10-AID pGAL-ndc10^trimer_E^*(SBY24687) strains were grown on either 2% glucose, 2% raffinose or 2% raffinose and galactose. 500 uM auxin was used to degrade the Ndc10-AID protein. c). Mtw1-3xmYPet and Spc110-mTurquoise2 label the kinetochores and spindle pole bodies in WT (SBY24595), *ndc10-AID, ndc10^trimer_E^* (SBY24691), *ndc10-AID, ndc10^trimer_A^* (SBY24692), *ndc10-AID, pGAL-ndc10^trimer_E^*(SBY24687), *and ndc10-AID, pGAL-ndc10^trimer_A^* (SBY24689) cells. The white arrowheads indicate strong kinetochore foci which were missing in mutant cells. Yellow arrowheads indicate chromosomes that segregated into two daughter cells. Representative pictures are shown for anaphase cells that were determined based on spindle pole body positions. The scale bar is 5 um. d). Quantification of kinetochore signal strength by measuring the Mtw1-3xmYPet in all cell cycle stages. All the images were normalized with the same threshold and the foci that are equally strong to WT Ndc10 are listed as strong while faint Mtw1 foci are listed as weak. When no foci were observed in a cell, it is listed as none. 200 cells were counted for each replicate, and three biological replicates were used to measure the significance using a two-tailed t test. **** P<0.00001, *** P< 0.0001. e). Quantification of chromosome segregation by observing DAPI signals. Only anaphase cells are counted, marked by separated Spc110-mTurquoise2 in daughter cells. When an anaphase cell has DAPI signals evenly distributed into the daughter cells, it is listed as segregation. When an anaphase cell has only one DAPI signal, it is listed as no segregation. ∼60 cells were counted for each replicate, and three biological replicates were used to measure the significance using a two-tailed t test. **** P<0.00001, *** P< 0.0001. f). Sequence alignment of Ndc10^D3^ through many fungal species. The residues colored in yellow are highly conserved and the residues colored in blue are medium conservation. The DNA binding residues are highlighted with a red box and the residues that are critical for trimer formation are colored in a green box. All trimer residues are conserved throughout the alignment. g). A hypothesized early kinetochore assembly stage. The models of CBF3 complex and CCAN were solved in this work. The nucleosome was used from PDB: 8ow0. h). Proposed intermediate states for kinetochore assembly related to g). The CBF3core is colored in green. The Ndc10^D1-D2^ and Ndc10^D4-D5^ are colored in yellow while the D3 trimer region is colored red. Cse4-H4 is colored in cyan. Scm3 is colored in dark blue. The unstructured region between D2-D3 and D3-D4 is illustrated with curved lines. It is not clear which or how many Ndc10 D4-D5 proteins recruit Scm3 to the kinetochore. Potential phosphorylation is colored in purple with letter P. Details of model are discussed in text.

To determine whether the mutants affect kinetochore assembly, we performed a bulk kinetochore assembly assay after auxin addition to the strains^38^. However, we noticed that the steady state levels of the Ndc10 mutants were lower than WT Ndc10 (Supplementary Fig. 5c). To control for this, we expressed the Ndc10 mutants from the galactose inducible promoter to restore their abundance prior to performing the assembly assay (Supplementary Fig. 5c). Under these conditions, there was a clear defect in kinetochore assembly in lysates from both trimer mutants (Supplementary Fig. 5d, lanes 4 and 6). Although the overexpressed Ndc10 proteins associated with centromeric DNA at levels comparable to wild type, recruitment of inner kinetochore components was strongly impaired and the cells remained inviable, indicating that Ndc10 trimerization is required for efficient inner kinetochore assembly rather than for initial CBF3–DNA binding (Supplementary Fig. 5d, lanes 4 and 6, Fig. 5b).

To determine the *in vivo* consequences, we expressed the *ndc10* mutants in an *ndc10-AID* strain with fluorescently marked kinetochores (Mtw1-3xmYPet)^49^ and spindle pole bodies (Spc110-mturquoise2)^50^. Cells were synchronized in G1 with α-factor in the presence of auxin and then released into the cell cycle with auxin to maintain Ndc10 depletion. After 105 mins, most cells had entered anaphase as defined by the separation of the two spindle pole bodies into each daughter cell (Fig. 5c). In WT cells, the kinetochore signals were strong and co-localized near the spindle pole body in both metaphase and anaphase cells. However, in *ndc10 ^trimer_A^* mutant cells, the Mtw1 signal was extremely weak. The phenotype was even more striking in the *ndc10 ^trimer_E^* mutant strain where most cells completely lost Mtw1 signal (Fig. 5c-d). We quantified chromosome segregation by DAPI staining and scoring cells with separated spindle pole bodies. In the wild-type strain, >90% of cells segregated chromosomal DNA equally, with a mild defect likely reflecting the presence of multiple epitope tags. In contrast, both *ndc10* trimer mutants exhibited severe chromosome segregation defects, with DNA remaining in the mother cell despite spindle elongation (Fig. 5c,e). The lack of kinetochore assembly explains why the cells do not arrest in metaphase via the spindle assembly checkpoint and are able to elongate their spindle but fail to segregate chromosomes. These observations, together with the biochemical assembly defects, indicate that Ndc10 trimerization is essential for robust kinetochore assembly *in vivo* and for subsequent chromosome segregation.

## Discussion

We developed an efficient system to purify native kinetochore complexes from budding yeast for cryoEM structural studies. This method complements reconstitution approaches that control stoichiometry and composition by enriching native kinetochore complexes that retain post-translational modifications and native CEN DNA. By assembling *de novo* kinetochores with native components, we were able to identify inner kinetochore complexes that had not been previously detected and speculate that some of the structures may be intermediate steps in kinetochore assembly. However, we did not detect outer kinetochore complexes, presumably because our method relied on immunoprecipitation of the inner kinetochore protein Ame1. In the future, performing an IP against outer kinetochore proteins after *de novo* assembly may identify additional features of the outer kinetochore.

Although CEN DNA was essential for *de novo* kinetochore assembly, we did not detect any clear DNA density in the CCAN dimer density map. We also did not visualize the largely unstructured Mif2 and Scm3 proteins that interact with both CCAN and DNA^51,52^. The requirement for CEN DNA in *de novo* assembly, together with the abundance of Mif2 and the presence of Scm3 in the assembly-IPs (Table S1), suggests that CEN DNA is present but unresolvable in these structures, likely because it is contacted primarily through these largely unstructured proteins. We similarly did not detect a Cse4 nucleosome in complex with any of the inner kinetochore structures, despite confirming its presence in the assembly-IP by immunoblotting and mass spectrometry. Therefore, even in the context of native kinetochore subcomplexes, the Cse4 nucleosome is not stable enough to be resolved at high resolution. One possibility is that the nucleosome-containing complexes are rare or structurally variable, which prevented us from obtaining an averaged density map. In the future, this difficulty might be overcome by enriching assembled kinetochores on grids coated with antibody^53^ or covalently decorating CEN DNA^54^ on the grid to perform assembly assay on grids.

Our assembly-IP protocol enriched kinetochore proteins via the CCAN component Ame1. However, the CBF3-CEN complex we purified isn’t known to bind to Ame1 directly. One explanation is that a larger CBF3-CEN-CCAN complex collapsed during vitrification, leaving only the CBF3-CEN portion resolved. To explore this, we carefully analyzed negative stain datasets of Ame1-3XFLAG assembly-IP material and generated larger 2D classes but did not observe any particles that resembled a stable CBF3-CEN-CCAN complex. An alternative explanation is that the CBF3-CEN and CCAN-containing complexes are linked only through CEN DNA, which may be partially unwrapped and highly flexible because the Cse4 nucleosome is not stable under our conditions. In that case, distinct regions of the same CEN DNA molecule could engage either CBF3–Ndc10 or Cbf1–CCAN, but the intervening DNA would not average into a continuous density, so we would solve structures corresponding to the CDEI-proximal and CDEIII-proximal ends separately. Future studies that use other inner kinetochore subunits as baits for assembly-IP, or that stabilize nucleosome-containing complexes, may help recover additional native assemblies.

We identified a previously unknown structure of Ndc10^D3^ that is essential for kinetochore assembly and cell viability. In contrast to a prior proposal that this region mediates Ndc10 dimerization^42^, our data show that D3 forms a trimeric assembly. Consistent with a conserved role for this interface, sequence alignment of multiple fungal Ndc10 orthologs reveals strong conservation of the hydrophobic residues that stabilize the trimer (Fig. 5f), and published fluorescent copy-number measurements of Ndc10 at kinetochores^55^ are compatible with a trimeric stoichiometry. Although Ndc10 is a kinetochore protein throughout the cell cycle, it also localizes to the spindle and bud neck after anaphase onset^31^. One speculative possibility, consistent with earlier models invoking an extended multimeric Ndc10 assembly over the centromere^30^, is that the third D3-associated Ndc10 copy could extend toward spindle microtubules while two Ndc10 molecules remain CBF3-bound. Because Ndc10 D3 also contacts CEN DNA in our structure, an important goal for future work will be to determine whether this interaction is sequence-specific and to define the consequences of disrupting it *in vivo*.

We also found that inclusion of the Ndc10 trimer correlates with stronger CBF3–DNA density in the native CBF3–CEN complex, and that native CEN sequences adopt a more sharply bent conformation than in prior reconstituted structures^21^. Given that the budding yeast centromere is a poor template for nucleosome assembly *in vitro*, these properties may help lower the energetic barrier to forming or stabilizing a Cse4 nucleosome on the AT-rich CEN DNA, a possibility that remains to be tested directly. Although it is unclear why Ndc10 trimerization was not observed in earlier reconstituted complexes, the differences we detect could reflect endogenous post-translational modifications or full-length protein contexts present in the native material. In support of this possibility, mass spectrometry of the assembly-IP identified several phosphorylation sites on Ndc10, and defining their functional contributions to Ndc10 trimerization and kinetochore assembly will be an important direction for future work.

Because native CEN DNA was used, our structures reveal two pronounced DNA bends, one near the CDEII–CDEIII junction and another between CDEI and CDEII, that were not apparent in previous reconstitutions using modified CDEII sequences. When we place a centromeric nucleosome (PDB: 8OW0) onto a model preserving these bends, there isn’t enough room for two copies of CCAN, two copies of CBF3 and the Ndc10 trimer without steric clashes. However, we can fit a model containing one CCAN, two CBF3 cores and an Ndc10^D1-D2^ (Fig. 5g). Based on this geometry, we propose that this structure and other native complexes we identified represent favored intermediates within an ensemble of early kinetochore assembly states (Fig. 5h). One possibility is that when CBF3 engages CDEIII and the surrounding pericentromeric DNA, it may form a dimeric CBF3–DNA assembly that is stabilized by the Ndc10 D3 trimer interacting with one Ndc10 D1–D2 domain. This interaction would sharpen the bend near CDEIII-CDEII junction to facilitate nucleosome formation when Ndc10 D4–D5 recruits Scm3–Cse4–H4^42^. Cbf1 and CCAN association with CDEI–proximal DNA and the emerging Cse4 nucleosome will help stabilize the nucleosome and one CCAN around the bent CDEII. As assembly proceeds, at least one Ndc10 copy disengages from the CBF3 core while remaining linked through the D3 trimer, consistent with the weak Ndc10–CBF3core interaction^26^ and Ndc10’s spindle localization later in mitosis^31^.

Although we and others detect a CCAN dimer *in vitro*, the Fiber-seq analysis shows that disrupting the interface between the Ndc10 D3 trimer and the Ndc10 D1/D2 region in the *ndc10^6A^* mutant specifically reduces protection over CDEIII, implicating CBF3 rather than a second copy of CCAN in this footprint. At present, we do not know whether the CCAN dimer we observe represents a physiological intermediate or an assembly state favored under our *in vitro* conditions. We also do not know if a second CBF3 complex on periCEN DNA is an experimental artifact, although our Fiber-seq data suggests there is likely a second copy of CBF3 instead of CCAN in this region.

In summary, we purified and identified native yeast high resolution kinetochore structures and identified conformations that were not previously detected in reconstitutions using recombinantly purified proteins. This work allowed us to elucidate a previously unknown Ndc10 trimerization that is conserved in sequence and essential for kinetochore assembly and chromosome segregation. Our work supports the previous proposal that Ndc10 is a central scaffold for inner kinetochore assembly^42^. Because many DNA-protein machines, such as chromatin remodelers and transcription complexes, assemble through dynamic, multi-step pathways, our results suggest that structural studies of other systems may similarly benefit from using natively assembled complexes.

## Materials and Methods

### Yeast strains

Media and genetic and microbial techniques were essentially as described^56^. Yeast strains used in this study are listed in Table S2. The 6His–3XFLAG epitope tagging of the endogenous *AME1, Mif2* and *CHL4* genes was performed using a PCR-based integration system using primers SB2434-SB2435 and plasmid pSB1590 as a template. All tagged strains we constructed are functional *in vivo* and do not cause any detectable growth defects or temperature sensitivity.

### Growth and lysate preparations from budding yeast

All yeast growth was performed as described previously^35^. Briefly, yeast were grown in YPD (1% yeast extract, 2% peptone, 2% D-glucose). Large cultures of SBY21782 (*AME1-3XFLAG*) were grown on shakers (220 rpm) at 23 °C. Cultures were treated with benomyl at a final concentration of 30 μg/ml (1:1 addition of 60 μg/ml benomyl YEP media) for 2 hours at 23 °C and then harvested by centrifugation for 10 minutes at 5000xg at 4 °C. Kinetochore material was purified based on a previously described protocol^35,38^. Briefly, the endogenous *AME1* kinetochore gene was C-terminally tagged with 6xHis and 3xFLAG. Harvested yeast were resuspended in Buffer H/0.15 (25 mM HEPES pH 8.0, 150 mM KCl, 2 mM MgCl_2_, 0.1 mM EDTA pH 8.0, 0.1% NP-40, 15% glycerol) supplemented with protease inhibitors, phosphatase inhibitors, and 2 mM DTT. After resuspension and re-spinning, yeast pellets were frozen in liquid nitrogen and lysed using a freezer mill (SPEX, Metuchen NJ). Lysates were first treated with 50 U/mL benzonase (Sigma-Aldrich) at room temperature for 30 minutes. The treated lysate was then clarified via tabletop centrifugation at 16,000g for 30 minutes and the supernatant was pooled. The sucrose gradient was generated by layering 1.5 mL of 40%, 25% and 10% sucrose in Buffer H/0.15 with no glycerol (25 mM HEPES pH 8.0, 150 mM KCl, 2 mM MgCl_2_, 0.1 mM EDTA pH 8.0, 0.1% NP-40) supplemented with protease inhibitors, phosphatase inhibitors and 2 mM DTT. The pooled supernatant was loaded on sucrose gradient. The lysate was further clarified via ultracentrifugation at 27,000 rpm for 16 hours, allowing contaminant proteins to pellet. The clarified layer was extracted with syringe. The extract was snap frozen with liquid nitrogen and stored in −80 °C until use.

### Purification of kinetochore complexes

For Ame1-3XFLAG IP, extracts prepared as described above were incubated with magnetic α-FLAG antibody conjugated Dynabeads (Invitrogen, Waltham MA) for 90 minutes at 4 °C with rotation. For Ame1-3XFLAG assembly IP, 18,750 ng sonicated single strand salmon sperm DNA (ssssDNA) per ml lysate was incubated with the extract on ice for 15 minutes and 625 ng 500 bp CEN DNA per ml lysate was rotated at 23 °C for 90 mins before adding magnetic α-FLAG antibody conjugated Dynabeads. The sequence of 500bp CEN DNA is:

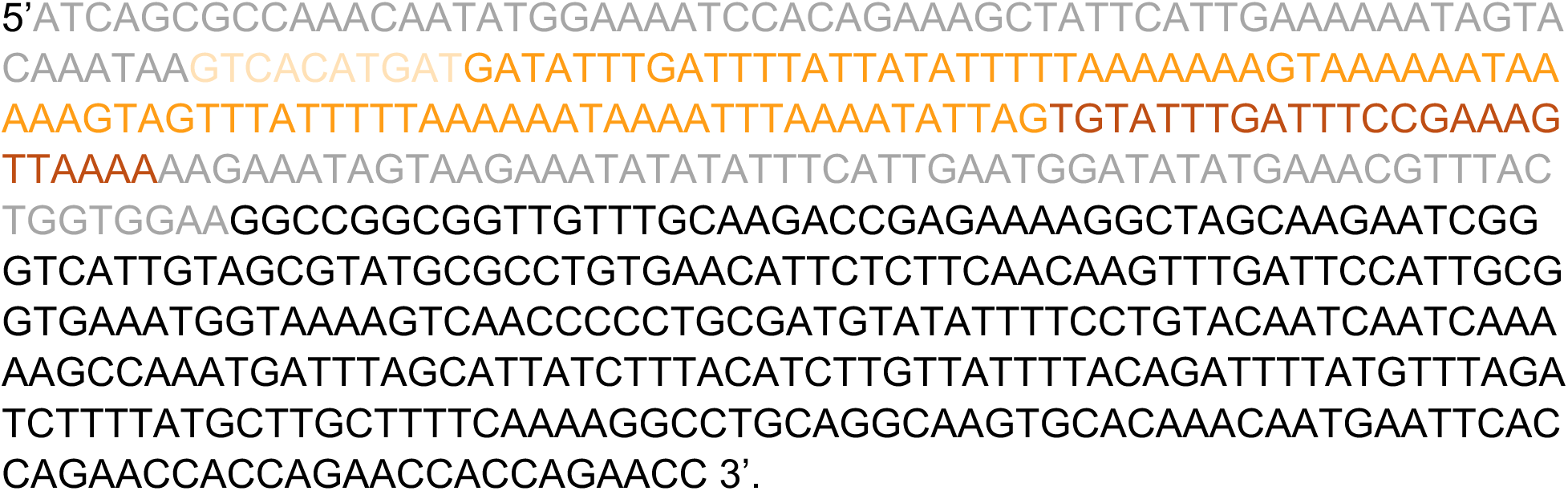

The periCEN nucleotides are colored in grey. The CDEI nucleotides are colored in light orange. The CDEII nucleotides are colored in orange. The CDEIII nucleotides are dark orange. The linker DNA is colored in black.

For immunoblotting, silver stain, mass spectrometry and cryoEM sample preparation, the Dynabeads were washed with 10x bead volumes of Buffer L/0.175 (25 mM HEPES pH 7.6, 175 mM KGlutamate, 6 mM Mg(OAc)_2_, 0.1 mM EDTA pH 7.6, 0.5 mM EGTA-KOH, pH7.6, 0.1% NP-40, 15% glycerol) 5 times. The last 3 washes omitted DTT and phosphatase inhibitors. For immunoblots and silver stain, kinetochores were eluted with 0.5 mg/ml 3xFLAG peptide in Buffer H/0.15 lacking DTT and phosphatase inhibitors. For mass spectrometry, kinetochores were eluted from Dynabeads with 0.2% RapiGest (Waters Corporation, Milford MA) in 50 mM HEPES pH 8.0. For negative stain electron microscopy and cryo-electron microscopy, kinetochores were washed with 10x bead volumes of Buffer H/0.15 5 times (the last 3 washes omitting DTT and phosphatase inhibitors), followed by one wash in Buffer L/0.175-EM (25 mM HEPES pH 7.6, 175 mM KGlutamate, 6 mM Mg(OAc)_2_, 0.1 mM EDTA pH 7.6, 0.5 mM EGTA-KOH, pH7.6) and eluted in 20uL 0.5 mg/ml 3xFLAG (Genscript, Piscataway NJ) peptide at room temperature for 30 minutes.

### Negative stain EM

Eluates from Ame1-3XFLAG IP or assembly-IP were used for negative stain electron microscopy. For both samples, 4 μL of eluate was diluted with Buffer L and deposited on 400 mesh continuous carbon grid (Electron Microscopy Sciences) for 1 minute separately. The grids were washed twice with Milli-Q water and stained with 0.75% uranyl formate (Electron Microscopy Sciences) for 1 minute. The EM grids were air dried before loading into Talos L120C (ThermoFisher). The data was collected at 1.991Å/pixel and processed in cryoSPARC^43^.

### Analysis of correlation between CEN DNA and CCAN dimer

Whole cell extracts were prepared as described above. The extract was equally separated into three parts and 0 ng/mL, 607 ng/mL or 1200 ng/mL of 500 bp CEN3 DNA was added to each part and incubated at room temperature for 90 minutes. The assembled kinetochores were purified as described in “purification of kinetochore complexes”. The three eluates were used to prepare three batches of negative stain grids and processed individually. Particles from 2D classifications that represent both apo-CCAN and CCAN dimer were counted as total CCAN. The ratio between CCAN dimer/ total CCAN was calculated.

### CryoEM grid preparation and data collection

For Ame1-3XFLAG IP eluate grid preparation, 3uL undiluted eluate was applied on 1.2/1.3 gold grids (Au 300, Electron Microscopy Science) covered freshly with graphene oxide (REF) and incubated for 30 s at 4 °C with 100% relative humidity and vitrified using a Vitrobot (Mark IV, Thermo Fisher). The grids were imaged under a Glacios transmission electron microscope (Thermo Fisher Scientific) operated at 200kV. The microscope is equipped with a Gatan K3 summit direct detection camera (Gatan, Pleasanton). A total of 3,821 movies were collected at a pixel size of 1.384 Å. For each micrograph, a 50-frame movie stack was collected with total exposure at 50 e^−^/Å^2^.

For Ame1-3XFLAG assembly IP eluate grid preparation, 1.2/1.3 holey titanium grids (Single Particles) were plasma cleaned (plasma cleaner brand) for 30 s. 3 μL undiluted Ame1-FLAG assembly-IP eluate was applied on grids and incubated for 30 s at 4 °C with 100% relative humidity and vitrified using a Vitrobot (Mark IV, Thermo Fisher). The frozen grids were imaged under a Titan Krios G3 transmission electron microscope (Thermo Fisher Scientific) operated at 300 kV. The microscope is equipped with a Gatan K3 summit direct detection camera (Gatan, Pleasanton). A total of 12,069 movies were collected at a pixel size of 1.07 Å. For each micrograph, a 50-frame movie stack was collected with total exposure at 50 e^−^/Å^2^.

### CryoEM data processing

The two sets of collected movies were motion-corrected and dose-weighted in cryoSPARC separately^43^. Contrast transfer function (CTF) corrected micrographs were used for blob picking and particles that belonged to well-resolved 2D classes were used for topaz picking^57^ for second round particle picking. For Ame1-3XFLAG IP eluate samples, these particles only include apo-CCAN complexes and the particles were cleaned through multiple rounds of 2D classification and heterogeneous refinement. A total of 88,088 particles was used for NU-refinement and a 3.9 Å density map was acquired. These particles include all four structures described in this paper. 744,531 particles were downsized and used for 2D classification. Classes that belong to apo-CCAN, Cbf1-CCAN-CEN, CCAN dimer, CBF3-CEN, and 40S, 60S and 80S ribosomes were used to run *ab initio* reconstructions, respectively. The remaining junk particles were also used to run ab initio reconstruction and separated into four classes. After obtaining the 10 initial models, we used them for the first round of heterogeneous refinement. Particles that belong to the 6 good classes were used for a second round of heterogeneous refinement using the 10 initial models. We ran heterogeneous refinement for 6 rounds until the particles that belong to 6 good classes didn’t change dramatically between runs and then we re-extracted selected particles at pixel size 1.07 Å and box size 424×424. We ran NU-refinement on each initial model (Supplementary Fig. 2). To improve local resolution, we did local refinement on the apo-CCAN, CCAN dimer and CBF3-CEN complexes. The resolution for each density map was assessed using the gold-standard criterion of Fourier shell correlation (FSC), with a cutoff at 0.143^58,59^, between 2 half maps from 2 independent half-sets of data (Supplementary Fig. 2).

### Model building

#### a). CCAN monomer

The initial model of apo-CCAN that was previously solved (PDB: 6QLE) was roughly fit into a density map that was solved in this work. The Ame1-Okp1 arm fit well while there was a misfit on the HIK arm. The HIK arm structure was fit into a density map and the extra density present in our structure was modeled using coot^60^. The new model was refined with Phenix^61^ and the Ramachandran outliers were fixed in coot again. This procedure was repeated until there were no outliers present in the structure.

#### b). Cbf1-CCAN-CEN

Cbf1 and CEN structures were predicted with AlphaFold 3^39^ and used as an initial model. The HIK head and Cnn1-Wip1 were modeled using 8OVW. The CCAN model from a) was fit into a density map. All the models mentioned above were combined and refined using Phenix^61^ once. The CEN DNA didn’t fit into our density map, so we modeled the DNA in coot^60^. A similar approach was taken that we refined iteratively with coot and Phenix^64^ until the Ramachandran outliers were fixed.

#### c). CCAN dimer

Since the CCAN dimer density was relatively low resolution, we couldn’t decipher side chains in the model. Thus, we did a rigid body fit of two CCAN dimer models into the density. The HIK head model was also fit into the density map.

#### d). CBF3-CEN

The CBF3 and CEN were modeled using AlphaFold 3^39^. Two copies of CBF3 were docked into chimeric density, which combined both focused refined maps. A similar approach was taken as described in a) and b).

To predict the extra density sequence, we utilized a focused refined map and set the fulcrum at the center of the extra density to optimize resolution in this region. The new local map was used as input in Model-Angelo^44^ with the command: model_angelo build_no_seq -v map.mrc -o output. The output model sequence was used for an NCBI blast search for a similar protein peptide. The predicted sequence was aligned with Ndc10(579-639), and the new model was built by replacing the predicted residues into Ndc10 residues manually. The extra density model was then refined using the approach mentioned above.

### Immunoblot and silver stain analyses

For immunoblot analysis, cell lysates were prepared as described above. Protein samples were separated using pre-cast 4-12% Bis Tris Protein Gels (Thermo-Fisher Scientific, Waltham MA) for sodium dodecyl sulfate-polyacrylamide gel electrophoresis (SDS-PAGE) in MES buffer pH 7.0 (50 mM MES, 50 mM Tris, 0.1% SDS, 1 mM EDTA). For immunoblotting, a 0.45 μm nitrocellulose membrane (BioRad, Hercules CA) was used to transfer proteins from polyacrylamide gels. The antibodies used for immunoblotting were custom generated by Genscript (Piscataway, NJ) against recombinant proteins that were expressed and purified from *Escherichia coli* and then injected into rabbits^25,46^. Genscript affinity purified the antibodies using the recombinant proteins and the resulting antibodies were used at the following dilutions: α-Mif2 used at 1:3000; α-Ctf19 used at 1:5000; α-Okp1 used at 1:5000; α-Ame1 used at 1:5000; α-Chl4 used at 1:5000; α-Mtw1 used at 1:5000; α-Ndc80 used at 1:5000; α-Spc105 used at 1:5000. Genscript services were also used to generate a FLAG antibody and V5 antibody which were used at 1:10,000^25^. The antibodies against Pgk1 and H2A were purchased from Invitrogen (4592560 and 39235) and used at 1:10,000 and 1:2000. α-Ndc10^38^ and α-Cse4^62^ were used at 1:5000 and 1:500. The secondary antibodies used were a sheep α-mouse antibody conjugated to horseradish peroxidase (HRP) (GE Life sciences, Marlborough MA) at a 1:10,000 dilution or a donkey α-rabbit antibody conjugated to HRP (GE Life sciences, Marlborough MA) at a 1:10,000 dilution. Antibodies were detected using the Super Signal West Dura Chemiluminescent Substrate (Thermo-Fisher Scientific, Waltham MA). For analysis by silver stain, the gels were stained with Silver Quest Staining Kit according to manufacturer’s instructions (Invitrogen, Waltham MA).

### Serial dilution assay

The designated *S. cerevisiae* strains were grown in YPD overnight. 1mL culture was spun down for each of the strains and the pellets were resuspended in 1mL YEP, respectively. Then the cell concentration was measured on a spectrophotometer (Bio-Rad), and cells were diluted to OD600 = 1.0. Next, serial dilutions (1:5) were made in water in a 96-well plate and the wells were then spotted onto YPD or YEP plates supplemented with 2% galactose and 2% raffinose. Plates were grown at the indicated temperatures for 1–3 days prior to imaging.

### Fiber-seq

In-house Hia5 preparation and yeast Fiber-seq were performed as described^25,47^. In brief, 10 ml cultures of wild-type (SBY3) and *ndc10^6A^* (SBY24368) yeast cells were grown in YPD medium to mid-log phase and collected by centrifugation. Cells were washed once with cold dH_2_O and resuspended in cold KPO4/Sorbitol buffer (1 M sorbitol, 50 mM potassium phosphate pH 7.5, 5 mM EDTA pH 8.0) supplemented with 0.167% β-mercaptoethanol. Spheroplasts were generated by adding Zymolyase T100 (0.15 μg/mL final concentration; Amsbio) and incubating at 23 °C for ∼15 min on a roller drum. Spheroplasts were pelleted at 160 g for 8 min at 4 °C, washed twice with cold 1 M Sorbitol, and resuspended in 58 uL of Buffer A (1 M Sorbitol, 15 mM Tris-HCl pH 8.0, 15 mM NaCl, 60 mM KCl, 1 mM EDTA pH 8.0, 0.5 mM EGTA pH 8.0, 0.5 mM Spermidine, 0.075% IGEPAL CA-630). Spheroplasts were treated with 1 μL of Hia5 MTase (200U) and 1.5 μL of 32 mM S-adenosylmethionine (NEB) for 10 min at 25 °C. The reaction was stopped by addition of 3 μL of 20% SDS (1% final concentration) and high molecular weight DNA was purified using the Promega Wizard® HMW DNA extraction kit (A2920). Circular consensus sequence reads were generated from raw PacBio subread files and processed as previously described^63^. Reads were aligned to the April 2011 *sacCer3* yeast reference genome, and nucleosomes were identified using default fibertools parameters^64^. To control for minor technical differences in the overall m6A methylation rate between each sample, reads were subsetted to normalize the per-read methylation distribution across samples (https://github.com/StergachisLab/match-distribution). Fibers overlapping with the center of each sixteen centromeres were extracted. The nucleosome density was calculated by counting the number of nucleosomes that overlap with each base pair of the region of interest divided by the number of fibers overlapping with that position. The averaged nucleosome density profiles across all sixteen centromeres are shown in Fig. 4d, and an individual profile for CEN3 is shown in Fig. 4e and 4g.

### Reconstituted Ndc10^D3-D5^ purification

Plasmids derived from pET28b were used for expressing WT or mutant Ndc10^D3-D5^ with an N-terminal 6xHIS tag. The plasmids were transformed into Rosetta2 competent cells and single colonies were inoculated in terrific broth with 25 ug/mL ampicillin to OD_600_=0.5. The cultures were then induced with 0.1 mM IPTG at 18 °C overnight before being spun down at 5000g for 10 mins. The cell pellet was resuspended in Buffer H/1.0 (25 mM HEPES pH 8.0, 1M KCl, 2 mM MgCl_2_, 0.1 mM EDTA pH 8.0, 0.1% NP-40, 15% glycerol) supplemented with 2 mM PMSF, EDTA-free protease cocktail and 10 mM imidazole. The cells were sonicated at 50% amplitude for 4 minutes with 1 second on and 1 second off. The supernatant was clarified by centrifugation at 13,000 g for 30 mins before applying to a 2 mL Talon cobalt resin and mixing for 1 hr in the cold room. The resin was then washed with 40 mL Buffer H/0.5 (25 mM HEPES pH 8.0, 500 mM KCl, 2 mM MgCl_2_, 0.1 mM EDTA pH 8.0, 0.1% NP-40, 15% glycerol, 20 mM imidazole) and eluted with 150 mM imidazole in Buffer H/0.15 (25 mM HEPES pH 8.0, 150 mM KCl, 2 mM MgCl_2_, 0.1 mM EDTA pH 8.0, 15% glycerol, 150 mM imidazole). The purified eluates were verified using SDS-PAGE (Supplementary Fig. 7a).

### Mass photometry

Mass photometry experiments were performed using the TwoMP system by Refeyn^65^. Movies were collected with Acquire2024 R1.1 and data was analyzed using Discover 2024 R1.0. The autofocus function was used to find the focus plane using 19 μl of cold modified buffer H 0.15 (25 mM HEPES pH 8.0, 150 mM KCl, 2 mM MgCl_2_, 0.1 mM EDTA pH 8.0, 15% glycerol) on uncoated glass slides (Refeyn). Thyroglobulin monomer and dimer peaks, conalbumin and aldolase were used as standards. Purified WT or mutant Ndc10^D3-D5^ fragments were diluted to 20 nM in cold modified Buffer H/0.15 and applied as droplet for measurement. Representative histograms of WT and mutants are shown in Supplementary Fig. 7b.

### TIRFM imaging and analysis

TIRFM assay to detect *de novo* kinetochore assembly at the single molecule level was performed as previously described^45,46^. Briefly, all images were captured using a Nikon TE-2000 inverted RING-TIRF microscope. Images were acquired at a resolution of 512 × 512 pixels with a pixel size of 0.11 μm/pixel at a readout speed of 10 MHz. Atto-647-labeled CEN DNAs were excited at 640 nm for 300 ms, GFP-tagged proteins at 488 nm for 200 ms. Images were analyzed with the CellProfiler (4.2.6) to assess colocalization and quantify signals between the DNA channel (647 nm) and GFP channel (488nm). Results were processed and visualized using FIJI (https://imagej.net/software/fij). For photobleaching assays, images in the 488 nm channel were acquired every 100 ms. The DNA-channel (647 nm) was only imaged for the initial frame throughout the movie. Photobleaching steps were analyzed using MATLAB to extract intensity traces of the 488 nm channel, and step count was performed using the same method as described^25,66^.

### Fluorescent microscopy of fixed yeast cells

WT or mutant *ndc10* strains with Mtw1-mYPet^49^ and Spc110-mTurquoise2^50^ were inoculated starting from OD_600_=0.4. To synchronize cells in G1 phase, 1ug/mL α factor was added to the culture and inoculated for 3 hours. 500 uM auxin was added 20 minutes before releasing cells to degrade Ndc10-AID. The cells were then washed two times with YEP containing 500 uM auxin and released for 105 mins before fixation. The cells were fixed with 3.7% formaldehyde at room temperature for 10 mins. To visualize the DNA, fixed cells were incubated with 1 ug/mL DAPI in 1.2M Sorbitol, 1% Triton, 0.1M potassium phosphate for 10 mins at room temperature. The stained cells were then pelleted and resuspended in 1.2M Sorbitol, 1% Triton, 0.1M potassium phosphate before imaging.

Fixed cell images were acquired on a Deltavision Ultra deconvolution high-resolution microscope equipped with a 60x/1.42 PlanApo N oil-immersion objective (Olympus). equipped with a 16-bit sCMOS detector. On both microscopes, cells were imaged in Z-stacks through the entire cell using 0.5 µm steps. All images were deconvolved using standard settings. All the images were projected for Z stack max intensity and normalized using FIJI. Cell stage was identified using Spc110-mTurquoise2 signals marking the two spindle pole bodies.

For Mtw1-YPet signal counting, only cells with visible spindle pole body signals were counted. The cells with an Mtw1 signal as strong as the WT strains were assigned as a strong signal. The cells with much weaker Mtw1 signal were assigned as a weak signal. The cells with only a diffuse Mtw1 signal were assigned as no signal. For all measured strains, 200 cells were counted and the experiment was repeated 3 times for a two-tailed t test to calculate the significant difference.

To quantify the DNA distribution in daughter cells, only anaphase cells with fully separated spindle pole bodies were counted. Cells with DAPI signals that were evenly distributed into daughter cells were counted as normal segregation while DAPI signals that were not evenly distributed were counted as chromosome missegregation. The experiments were repeated three times on three batches of cells.

## Data availability

Mass spectrometry data generated in this study is available through Mass Spectrometry Interactive Virtual Environment (MassIVE, University of California San Diego) with the link https://massive.ucsd.edu/ProteoSAFe/dataset.jsp?task=a50a708407634efc951e1828867bd934. The density map and model of apo-CCAN are accessible with code EMD-75131 from EM Databank and 10FI from Protein Data Bank. The density map and model of CCAN dimer are accessible with code EMD-75213 from EM Databank and 10JC from Protein Data Bank. The density map and model of Cbf1-CCAN-CEN are accessible with code EMD-75095 from EM Databank and 10DQ from Protein Data Bank. The density map and model of CBF3-CEN are accessible with code EMD-75107 from EM Databank and 10EH from Protein Data Bank. The plasmids used in this study can be provided when requested. The Fiber-seq sequencing data generated in this study for the ndc10-6A mutant have been deposited in the NCBI Sequence Read Archive (SRA) database under the accession number PRJNA1427749. Fiber-seq data for the WT strain are available at SRA under accession number PRJNA1189155^25^.

## Acknowledgments

We are grateful to the Biggins and Asbury labs for critical reading of the manuscript and fruitful discussions. We thank Chip Asbury and Barry Stoddard for insightful comments on the manuscript. We thank the Proteomics & Metabolics shared resource facility at Fred Hutchinson Cancer Center for mass spectrometry sample processing and data analysis as well as the Fred Hutch Electron Microscopy & CryoEM Core for grid preparation, sample screening, data acquisition, and professional suggestions. We thank the HHMI CryoEM Shared Resource at Janelia Research Campus for cryoEM data acquisition. We also thank the Cellular Imaging team, especially Lena Schroeder and Hoku West-Foyle, for all their help. We are grateful to Alex Kaiser for his help with mass photometry and to Shane Neph, Yizi Mao, and Ben Mallory for their assistance with purifying the Hia5 enzyme, data analysis and sequencing library construction.

## Contributions

Conceptualization (M. Jiang and S. Biggins), data curation (M. Jiang, C. Hu, S. Hedouin and S. Biggins), Data acquisition (M. Jiang, C. Hu, S. Hedouin, A. Latino), model building (M. Jiang and Y. Arimura), Data analysis (M. Jiang, S. Hedouin, C. Hu, and S. Biggins), funding acquisition (S. Biggins, A. Stergachis, C. Hu), investigation (M. Jiang, C. Hu, S. Hedouin, A. Latino), project administration (S. Biggins, A. Stergachis), supervision (S. Biggins, A. Stergachis), validation (M. Jiang and S. Biggins), visualization (M. Jiang and S. Biggins), writing—original draft (M. Jiang and S. Biggins), writing—review and editing (M. Jiang, C. Asbury, B. Stoddard, Y. Arimura, A. Stergachis, S. Biggins).

## Author information

Mengqiu Jiang, Basic Sciences Division, Howard Hughes Medical Institute, Fred Hutchinson Cancer Center, Seattle, WA 98109, United States.

Changkun Hu, Basic Sciences Division, Howard Hughes Medical Institute, Fred Hutchinson Cancer Center, Seattle, WA 98109, United States.

Sabrine Hedouin, Basic Science Division, Fred Hutchinson Cancer Center, Seattle, WA, 98109, United States.

Angelica Andrade Latino, Basic Science Division, Fred Hutchinson Cancer Center, Seattle, WA, 98109, United States.

Yasuhiro Arimura, Basic Science Division, Fred Hutchinson Cancer Center, Seattle, WA, 98109, United States.

Andrew B. Stergachis, Division of Medical Genetics, Department of Medicine, Department of Genome Sciences, University of Washington, Seattle, WA, 98195, United States.

Sue Biggins, Basic Sciences Division, Howard Hughes Medical Institute, Fred Hutchinson Cancer Center, Seattle, WA 98109, United States.

## Ethics declarations

None declared.

