## Supplementary Data for "Native yeast kinetochore structures identify an essential inner kinetochore interaction"

**Supplementary Material**

1. Supplementary Figures
2. Supplementary Tables
3. Movie
4. References

Supplementary Figure 1

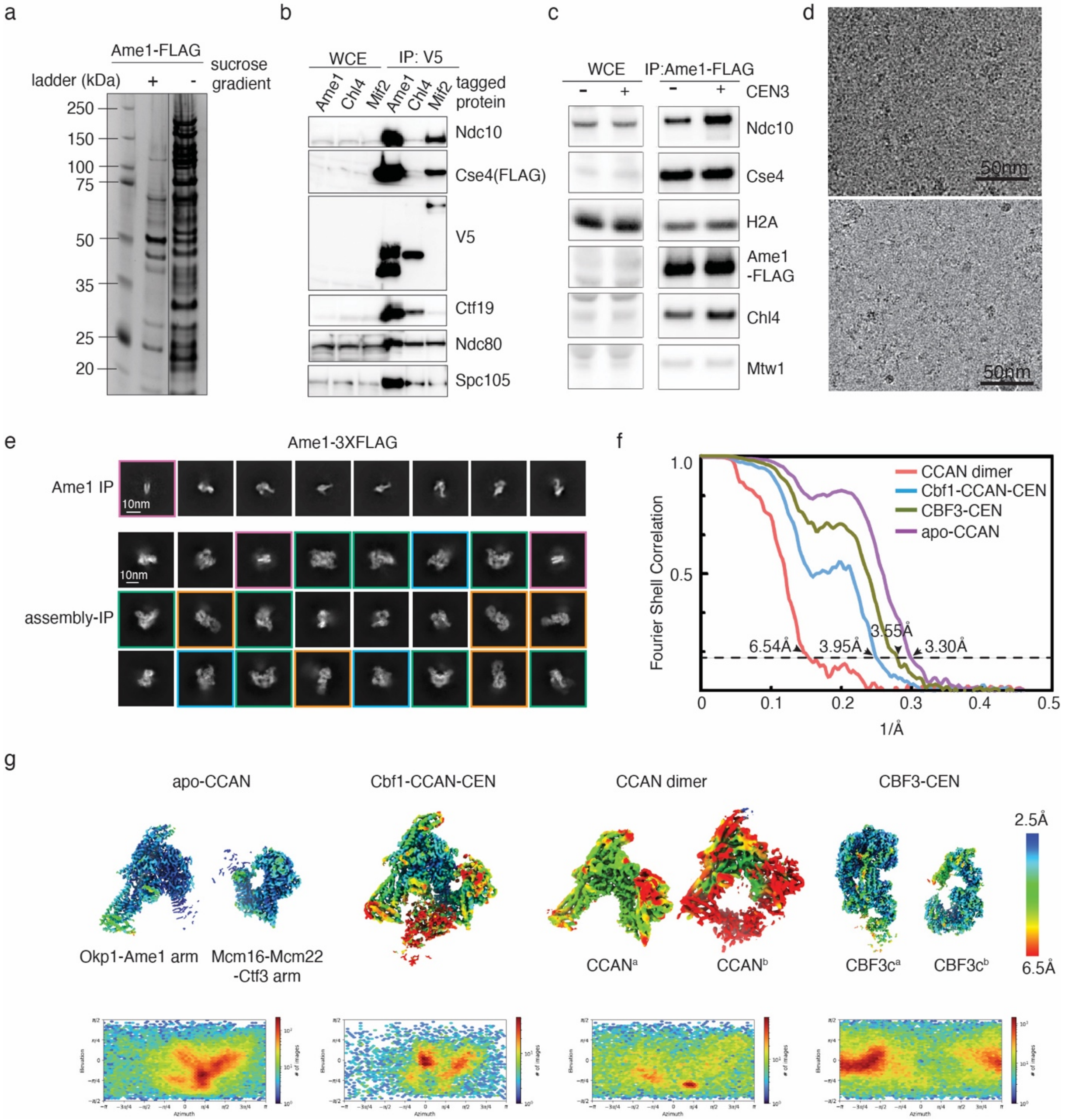


**Supplementary Figure 1. Examination of eluates from Ame1-3XFLAG IP or assembly-IP using sucrose gradient precleared lysates.**

a). Silver-stained analysis of SDS-PAGE on eluates from Ame1-3XFLAG (SBY21782) IP using precleared (sucrose gradient) or regular lysate. The two lanes were from the same gel but trimmed to remove the non-related lanes indicated by the black line separation.

b). Immunoblot of immunoprecipitation from lysates that had different kinetochore protein tags (SBY21489*: CHL4-3xV5,* SBY21488*: MIF2-3xV5,* SBY21120*: AME1-*

*3xV5*). The tagged protein is indicated, and multiple kinetochore proteins were detected by immunoblotting with the corresponding antibodies. WCE is whole cell lysate.

c). Immunoblots show enrichment of kinetochore proteins from Ame1 IP or assembly IP using antibodies against the indicated kinetochore proteins. WCE is whole cell lysate.

d). A representative micrograph of Ame1-3XFLAG IP eluate (upper panel) and assembly-IP eluate (lower panel).

e). 2D classification of Ame1-3XFLAG IP eluate (top row) or Ame1-3XFLAG assembly-IP eluate (bottom row). The averages of CCAN dimer are colored in green; the averages of Cbf1-CCAN-CEN are colored in blue; the averages of CBF3-CEN are colored in orange; the averages of nucleosome are colored in pink. The rest of averages are apo-CCAN.

f). Overall resolutions were estimated with Fourier shell correlation (FSC) at 0.143 standard[1]. The local resolutions are shown by coloring densities with rainbow colors, with highest resolution at 2.5 Å colored in blue and the lowest resolution at 6.5 Å and colored in red. The name for each density is labeled.

g). Local resolution estimation of the four structures solved. Blue color represents higher resolution and red color represents lower resolution. Angular distributions for each complex were shown in the bottom panel.

Supplementary Figure 2

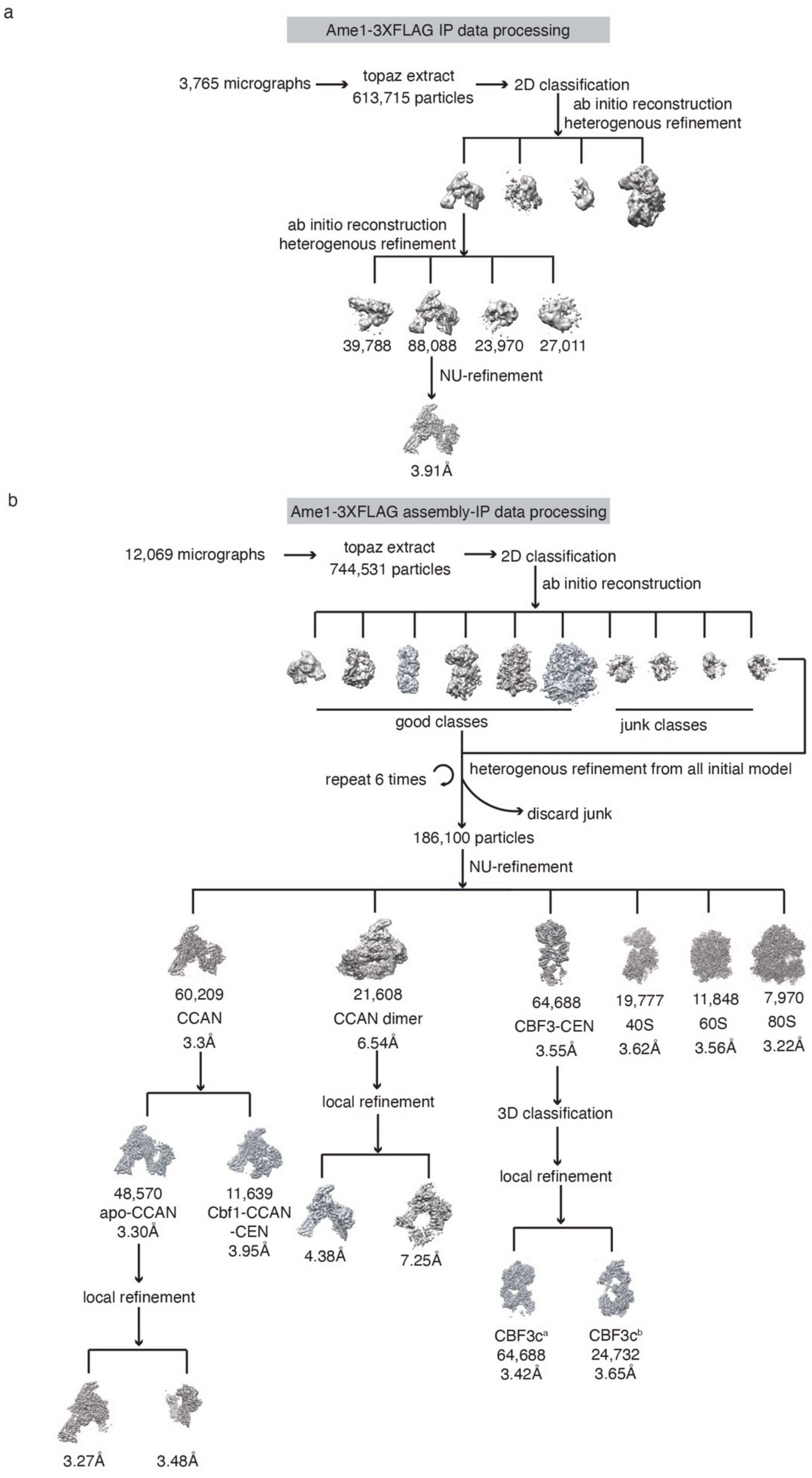

**Supplementary Figure 2. Flowchart of cryoEM data processing and overall resolution and local resolution of the structures identified from this work.**

a). Data processing of Ame1-3XFLAG IP eluate.

b). Data processing of Ame1-3XFLAG assembly-IP eluate.

Supplementary Figure 3

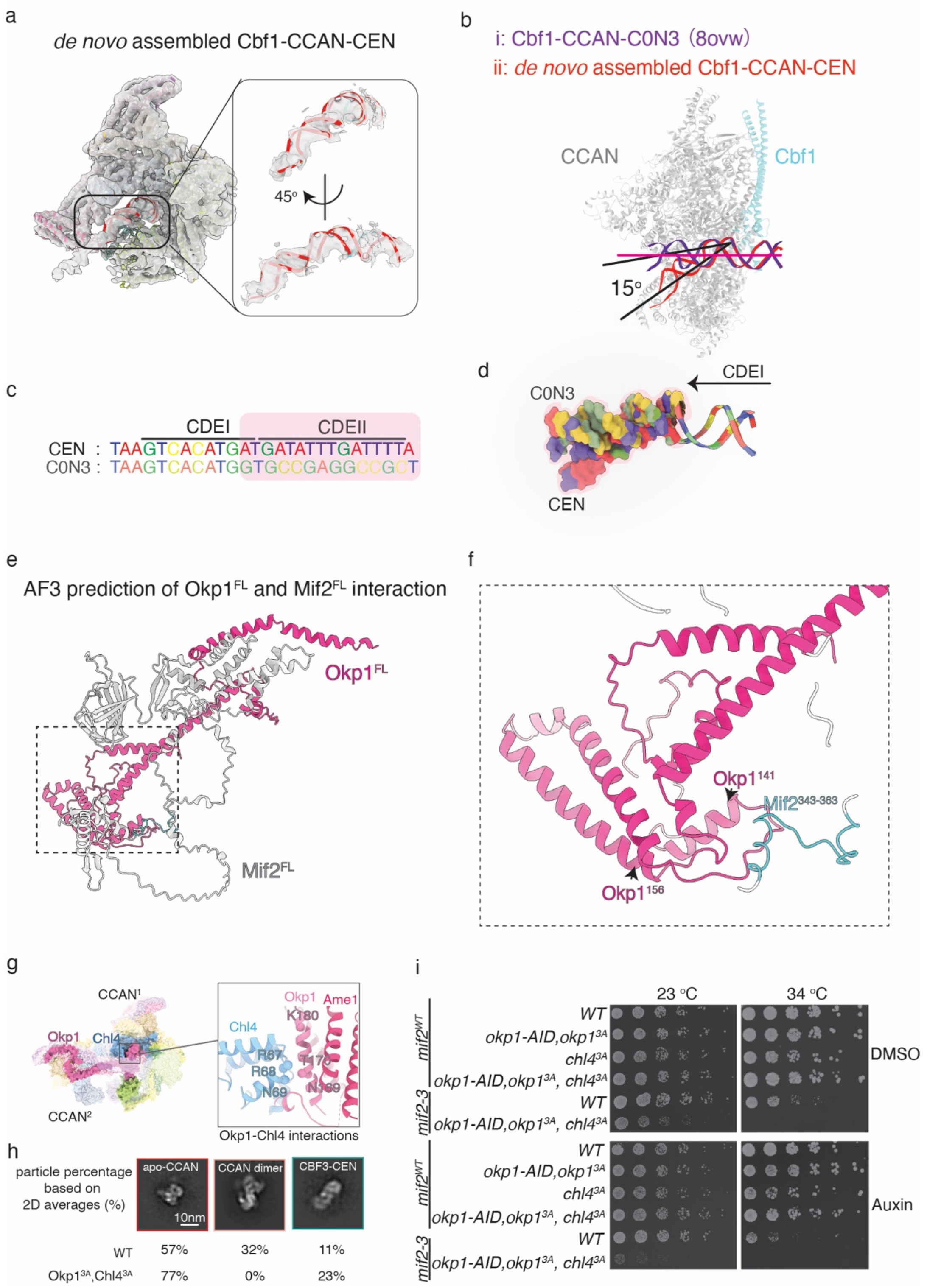


**Supplementary Figure 3. Comparison between native CCAN and recombinant reconstituted CCAN and DNA bending in the Cbf1-CCAN-CEN complex.**

a). Model of *de novo* assembled Cbf1-CCAN-CEN fitted in low pass filtered density map to show the trace of DNA. The black box area is enlarged and cropped density of CEN DNA is shown. Two views were provided to show the DNA groove density.

b). Superimposition of the native Cbf1-CCAN-CEN and the reconstituted Cbf1-CCAN-C0N3 (8OVW)[3]. The model of CCAN is colored in transparent grey and Cbf1 is colored in cyan. CEN DNA is red and C0N3 DNA is purple. The angle difference between the two DNA molecules is demonstrated by a $15^{o}$ angle.

c). Sequences of CEN and C0N3 DNA that interact with the CCAN-Cbf1 complex. The CDEI and part of the CDEII sequences are labeled. The sequence differences between CDEII are highlighted by transparent pink shade.

d). The two DNA sequences were superimposed with C0N3 transparent and CEN3 solid. The direction of DNA is labeled with arrows.

e). One AlphaFold model of proposed full-length (FL) Mif2 and Okp1 interaction. Okp1 is colored in hot pink and Mif2 is colored in grey.

f). Zoom-in view on the right, $Mif2^{343\text{-}363}$ fragment is colored in cyan.

g). Cartoon shows predicted interaction surface of CCAN dimer based on our EM density. Cα of residues that are involved in Okp1-Chl4 interface and Okp1-Ctf3 interface are shown.

h). Particle percentage of apo-CCAN, CCAN dimer and CBF3-CEN calculated from 2D classification using negative stain EM on WT or $okp1^{3A}chl4^{3A}$ mutant. In $okp1^{3A}chl4^{3A}$ mutant, no CCAN dimer is observed.

i). Viability assay using mutants on chl4 and okp1 under $23^{o}C$ or $34^{o}C$ are shown.

Supplementary Figure 4

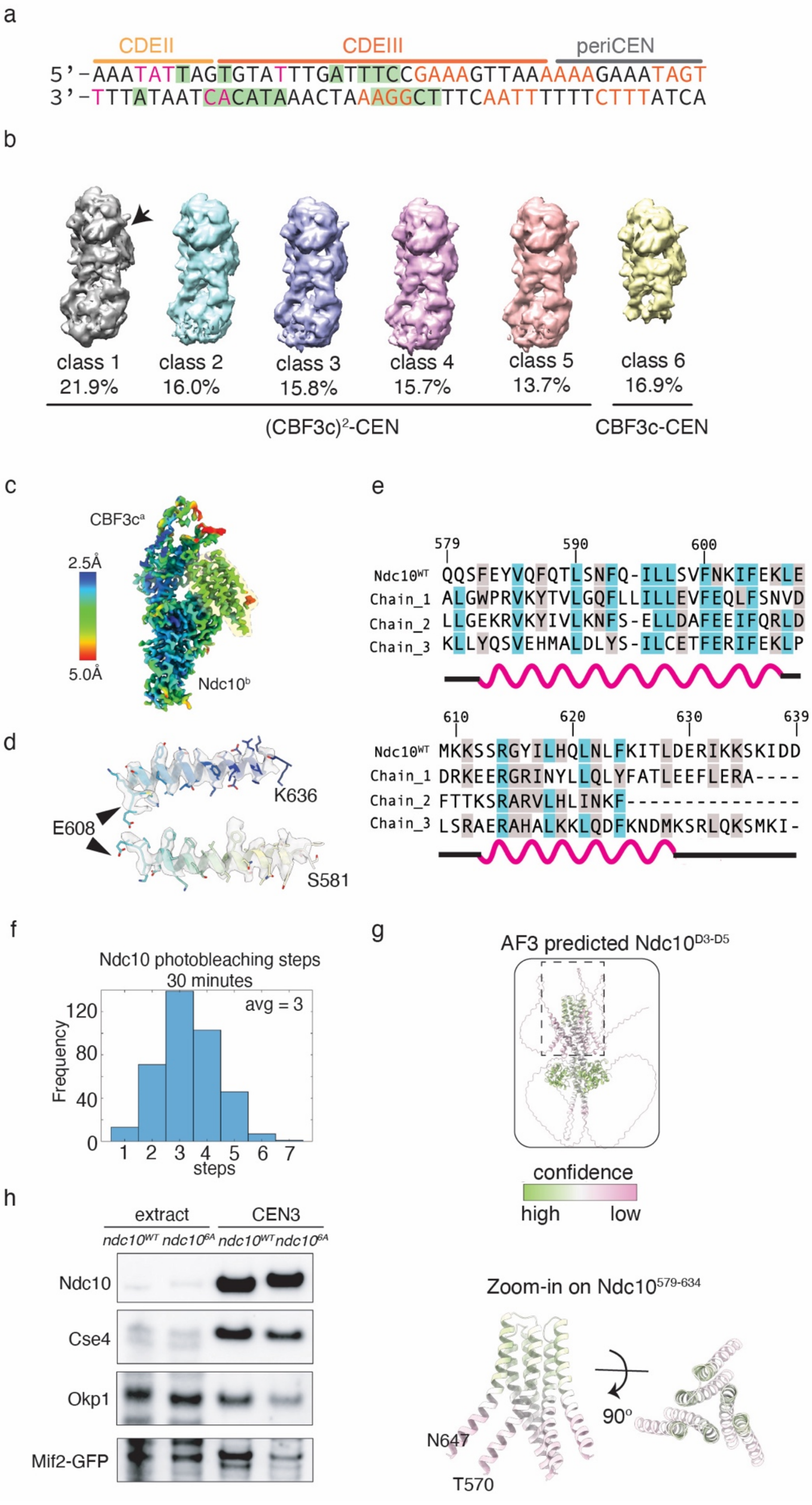

**Supplementary Figure 4. Interaction between CBF3c and CEN3 DNA.**

a). The regions of CDEII, CDEIII and periCEN DNA that are protected by CBF3c are labeled. Base pairs that interact with Ndc10 and extra density are colored in orange and pink, respectively. Base pairs that interact with Cep3 are included in green shaded boxes.

b). 3D classification of all 6 classes of CBF3c-DNA particles. The arrow indicates the extra density. The ratio of each class is labeled. Class 6 is a CBF3c monomer-DNA complex.

c). Local resolution of focus refined density map. The high-resolution density is blue and the low-resolution density is red. The color range is 2.5 Å to 5 Å.

d). Demonstration of model fitting into extra density with WT-Ndc10 sequence. The bulky side chain fitting is shown.

e). Sequence alignment for the predicted three chains aligned with WT-Ndc10. The highly conserved residues are blue, and the less conserved residues are grey. The secondary structure is shown in a cartoon.

f). The frequency of photobleaching steps on kinetochores assembled from extracts from Ndc10-GFP cells (SBY22191) via a single molecule TIRF assay after *de novo* assembly for 30 minutes.

g). AlphaFold 3 prediction of an Ndc10$^{D3\text{-}D5}$ trimer. The Ndc10$^{D3_trimer}$ is zoomed in and other regions were hidden.

h). Immunoblot of a kinetochore assembly assay on WT (SBY12796) or mutant *ndc10* (SBY23670) lysates.

Supplementary Figure 5

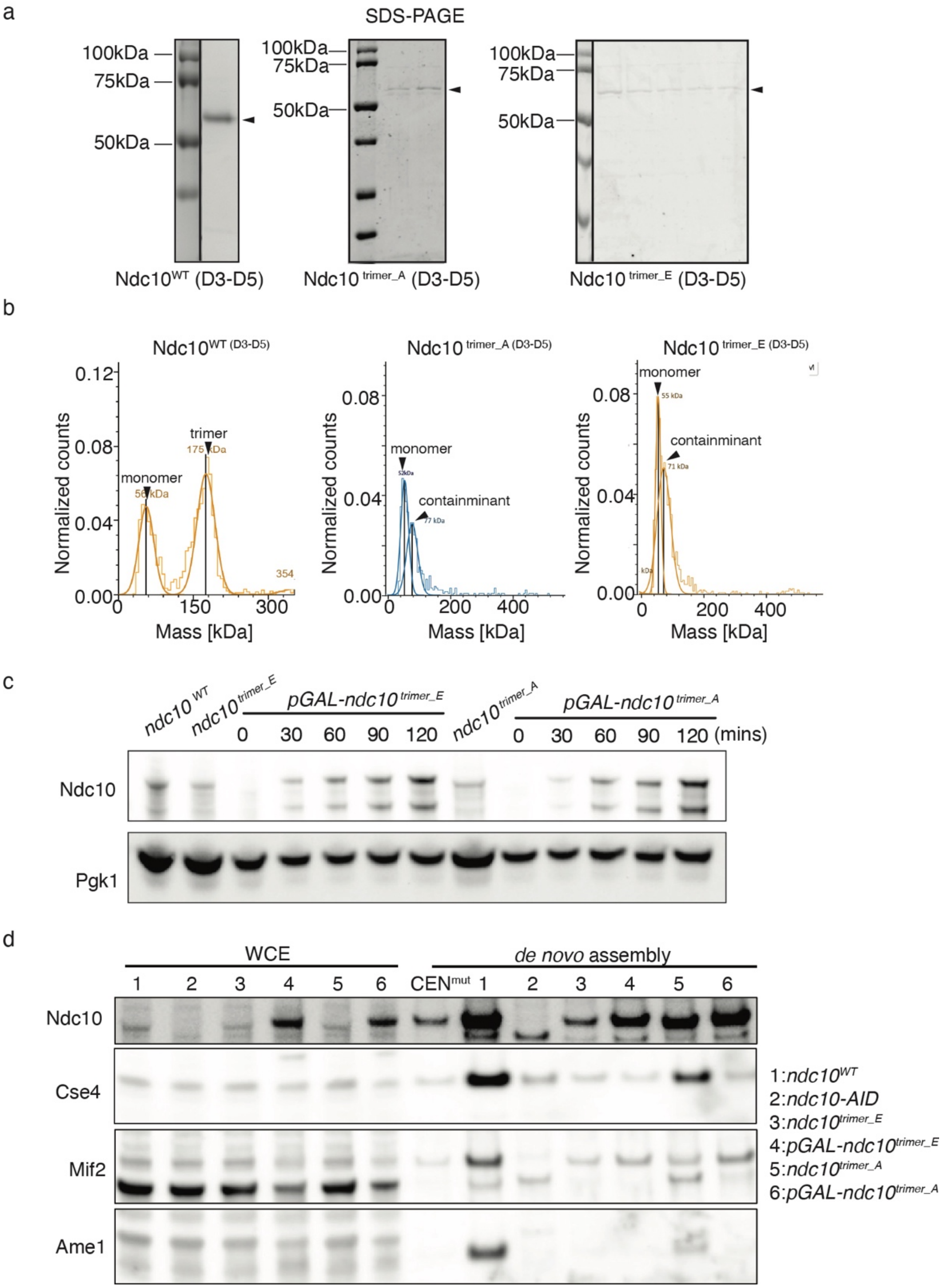


**Supplementary Figure 5. Characterization of ndc10 trimer proteins.**

a). SDS-PAGE of purified WT-Ndc10, Ndc10 $^{trimer\_E}$ and Ndc10 $^{trimer\_A}$ proteins. The lanes in each box were from the same gel but trimmed to remove the non-related lanes indicated by a black line separation.

b). Mass photometry measurements of purified Ndc10(D3-D5)$^{trimer\_A}$ and Ndc10(D3-D5)$^{trimer\_E}$. The peak at ~55 kDa is an Ndc10 monomer and the peak at ~70 kDa is a contaminant protein.

c). Immunoblot on whole cell extracts made from WT (SBY12796), *ndc10-AID* (SBY21327), *ndc10-AID ndc10$^{trimer_A}$* (SBY24533), *ndc10-AID ndc10 $^{trimer\_E}$* (SBY24478), *ndc10-AID pGAL-ndc10$^{WT}$* (SBY24690), *ndc10-AID pGAL-ndc10$^{trimer_A}$* (SBY24689), and *ndc10-AID pGAL-ndc10$^{trimer_E}$* (SBY24687) strains. Mutant protein overexpression was induced with 2% galactose and samples removed every 30 minutes. The overexpression was observed starting at 30 minutes. The loading control is Pgk1.

d). Immunoblot of kinetochore assembly assay on WT (SBY24587), *ndc10-AID* (SBY21327), *ndc10-AID ndc10$^{trimer_A}$* (SBY24533), *ndc10-AID ndc10 $^{trimer\_E}$* (SBY24478), *ndc10-AID pGAL-ndc10$^{WT}$* (SBY24690), *ndc10-AID pGAL-ndc10$^{trimer_A}$* (SBY24689), *ndc10-AID pGAL-ndc10$^{trimer_E}$* (SBY24687) extracts using antibodies against the indicated inner kinetochore protein.

**Table S1. Mass spectrometry result of Ame1-3XFLAG assembly-IP eluate.**

| Complex | Protein name | #PSMs |
|---|---|---|
| CBF3 | Ctf13 | 12 |
| | Cep3 | 24 |
| | Ndc10 | 50 |
| | Skp1 | 5 |
| CCAN | Okp1 | 219 |
| | Ctf19 | 80 |
| | Ame1 | 112 |
| | Mcm21 | 80 |
| | Nkp1 | 140 |
| | Nkp2 | 60 |
| | Ctf3 | 8 |
| | Mcm16 | 1 |
| | Mcm22 | 11 |
| | Chl4 | 19 |
| | Iml3 | 38 |
| | Cnn1 | 5 |
| | Wip1 | 1 |
| | Mif2 | 184 |
| Chaperone | Scm3 | 4 |
| Dam1c | Spc19 | n.d. |
| | Dad1 | 3 |
| | Spc34 | n.d |
| | Dad2 | n.d. |
| | Dad3 | n.d. |
| | Dad4 | n.d. |
| | Hsk3 | n.d. |
| | Dam1 | 7 |
| | Duo1 | 3 |
| | Ask1 | 2 |
| Kinase | Cdc5 | 14 |
| | Mps1 | 63 |
| | Pgk1 | 6 |
| Knl1c | Kre28 | 5 |
| | Spc105 | 10 |
| Ndc80c | Nuf2 | 28 |
| | Spc25 | 29 |
| | Ndc80 | 36 |
| | Spc24 | 39 |
| Nucleosome | H2A | 25 |
| | H2B | 31 |
| | Cse4 | 94 |
| | H4 | 38 |

#PSMs: Peptide Spectrum Matches

**Table S2. Strains used in this article**

**All strains are in the w303 strain background.**

| **Strain name** | **genotype** | **Used in Figure** |
|---|---|---|
| SBY3 | *MATa, bar1-1 ade2-1 can1-100 his3-11 ura3-1 trp1-1 leu2-3* | Fig.4 |
| SBY12796 | *MATa NDC10-3XFLAG:KAN bar1-1 ade2-1 can1-100 his3-11 ura3-1 leu-2-3* | Fig. 5, Supplementary Fig. 4-5 |
| SBY21120 | *MATa AME1-3XV5:KAN LEU2-3::pGAL-SCM3-mCherry:LEU2 URA3-1::CSE4-FLAG:URA3 CSE4^:KAN ade2-1 bar1^ trp1-1 his3-11 can1-100* | Supplementary Fig. 1 |
| SBY21327 | *MATα NDC10-3XV5-IAA7:KAN LEU2-3:: pGPD1-OsTIR1:LEU2 bar1-1 ade2-1 can1-100 his3-11 trp1-1 ura3-1* | Fig. 5, Supplementary Fig.5 |
| SBY21479 | *MATa mif2-3 ura3-1 leu2-3 ade2-1 can1-100 his3-11 trp1-1 bar1-1* | Fig. 4 |
| SBY21488 | *MATα MIF2-3XV5:HIS3 LEU2-3::pGAL-SCM3-mCherry:LEU2 CSE4-FLAG:URA3 CSE4^:KAN ade2-1 bar1^ trp1-1 can1-100* | Supplementary Fig. 1 |
| SBY21489 | *MATα CHL4-3XV5:HIS3 LEU2-3::pGAL-SCM3-mCherry:LEU2 CSE4-FLAG:URA3 CSE4^:KAN ade2-1 bar1^ trp1-1 can1-100* | Supplementary Fig. 1 |
| SBY21782 | *MATα AME1-3XFLAG:TRP1 ura3-1 his3-11 trp1-1 leu2-3 ade2-1 can1-100 bar1-1* | Fig. 1, Supplementary Fig.1 |
| SBY22094 | *MATα MIF2-GFP:KAN bar1-1 ade2-1 can1-100 trp1-1 his3-1 leu2-3 ura3-1* | Fig. 4 |
| SBY22191 | *MATα Ndc10-GFP:KAN bar1-1 ade2-1 can1-100 his3-11 ura3-1* | Fig. 3, Supplementary Fig. 4 |
| SBY23670 | *MATa ndc10-S493A-D494A-S497A-D501A-H505A-K507A-3xFLAG:KAN bar1-1 ade2-1 can1-100 trp1-1 his3-11 leu2-3 ura3-1* | Fig. 4, Supplementary Fig. 4 |
| SBY23913 | *MATα mif2-3 ndc10-S493A-D494A-S497A-D501A-H505A-K507A-3xFLAG:KAN bar1-1 ade2-1 can1-100 trp1-1 his3-11 leu2-3 ura3-1* | Fig. 4 |

| | | |
|---|---|---|
| SBY24238 | *MATα MIF2-GFP:KAN ndc10-S493A-D494A-S497A-D501A-H505A-K507A-3xFLAG:KAN bar1-1 ade2-1 can1-100 trp1-1 his3-1 leu2-3 ura3-1* | Fig. 4 |
| SBY24478 | *MATα TRP1-1::ndc10-F593E-L596E-F600E-F604E-3XV5:TRP1 NDC10-3XV5-IAA7:KAN LEU2-3::pGPD1-OsTIR1:LEU2 bar1-1 ade2-1 can1-100 his3-11 ura3-1* | Fig. 5, Supplementary Fig.5 |
| SBY24533 | *MATα TRP1-1::ndc10-F593A-L596A-F600A-F604A-3XV5:TRP1 NDC10-3XV5-IAA7:KAN LEU2-3::pGPD1-OsTIR1:LEU2 bar1-1 ade2-1 can1-100 his3-11 ura3-1* | Fig. 5, Supplementary Fig.5 |
| SBY24587 | *MATα TRP1-1::NDC10-3XV5:TRP1 NDC10-3XV5-IAA7:KAN LEU2-3::pGPD1-OsTIR1:LEU2 bar1-1 ade2-1 can1-100 his3-11 ura3-1* | supplementary Fig.5 |
| SBY24595 | *MATa TRP1-1::NDC10-3XV5:TRP1 NDC10-3XV5-IAA7:KAN LEU2-3::pGPD1-OsTIR1:LEU2 MTW1-mYPet:URA3 SPC110-mTurquoise2:TRP1 bar1-1 ade2-1 can1-100 his3-11* | Fig. 5 |
| SBY24687 | *MATα TRP1-1::pGAL-ndc10-F593E-L596E-F600E-F604E-3XV5:TRP1 NDC10-3XV5-IAA7:KAN LEU2-3:: pGPD1-OsTIR1:LEU2 bar1-1 ade2-1 can1-100 his3-11 trp1-1 ura3-1* | Fig. 5, Supplementary Fig.5 |
| SBY24689 | *MATα TRP1-1::pGAL-ndc10-F593A-L596A-F600A-F604A-3XV5:TRP1 NDC10-3XV5-IAA7:KAN LEU2-3::pGPD1-OsTIR1:LEU2 bar1-1 ade2-1 can1-100 his3-11 trp1-1 ura3-1* | Fig. 5, Supplementary Fig.5 |
| SBY24690 | *MATa TRP1-1::pGAL-NDC10-3XV5:TRP1 bar1-1 ade2-1 can1-100 his3-11 trp1-1 ura3-1* | Fig. 5, Supplementary Fig.5 |
| SBY24691 | *MATa TRP1-1::ndc10-F593E-L596E-F600E-F604E-3XV5:TRP1 NDC10-3XV5-IAA7:KAN LEU2-3::pGPD1-OsTIR1:LEU2 MTW1-mYPet:URA3 SPC110-mTurquoise2:TRP1 bar1-1 ade2-1 can1-100 his3-11* | Fig. 5 |
| SBY24692 | *MATa TRP1-1::ndc10-F593A-L596A-F600A-F604A-3XV5:TRP1 NDC10-3XV5-IAA7:KAN LEU2-3::pGPD1-OsTIR1:LEU2 MTW1-mYPet:URA3 SPC110-mTurquoise2:TRP1 bar1-1 ade2-1 can1-100 his3-11* | Fig. 5 |

**Table S3. Data collection and refinement statistics**

| | apo-CCAN (EMD-75131) (PDB: 10FI) | Cbf1-CCAN-CEN (EMD-75095) (PDB: 10DQ) | CCAN dimer (EMD-75213) (PDB: 10JC) | CBF3-CEN (EMD-75107) (PDB: 10EH) |
|---|---|---|---|---|
| **Data collection and processing** | | | | |
| Magnification | 64,000 | 64,000 | 64,000 | 64,000 |
| Voltage (kV) | 300 | 300 | 300 | 300 |
| Electron exposure (e–/Å$^2$) | 50 | 50 | 50 | 50 |
| Defocus range (μm) | -1~-2 | -1~-2 | -1~-2 | -1~-2 |
| Pixel size (Å) | 1.07 | 1.07 | 1.07 | 1.07 |
| Symmetry imposed | C1 | C1 | C1 | C1 |
| Final particle mages (no.) | 48,570 | 11,639 | 21,608 | 64,688 |
| Map resolution (Å) FSC threshold | 3.30 | 3.95 | 6.54 | 3.55 |
| **Refinement** | | | | |
| Model resolution (Å) FSC threshold | 3.30 | 3.95 | 6.54 | 3.55 |
| Model composition | | | | |
| Non-hydrogen atoms | 18835 | 25868 | 47029 | 40188 |
| Protein residues | 2458 | 3190 | 5820 | 4611 |
| DNA residues | 0 | 54 | 0 | 88 |
| R.m.s. deviations | | | | |
| Bond lengths (Å) | 0.004 | 0.003 | 0.004 | 0.008 |
| Bond angles (°) | 0.615 | 0.658 | 0.607 | 0.607 |

| | | | | |
|---|---|---|---|---|
| Validation | | | | |
| MolProbity score | 2.18 | 2.22 | 1.96 | 2.45 |
| Clashscore | 10.23 | 12.19 | 13.57 | 12.78 |
| Poor rotamers (%) | 2.51 | 2.95 | 0.00 | 4.05 |
| Ramachandran plot | | | | |
| Favored (%) | 95.04 | 96.14 | 95.35 | 94.63 |
| Allowed (%) | 4.75 | 3.83 | 4.61 | 5.19 |
| Disallowed (%) | 0.21 | 0.03 | 0.04 | 0.18 |

**Movie 1. Movement of CEN DNA when comparing native and reconstituted CBF3-CEN.**
